# Hide and seek: *de novo* identification in sugar beet reveals impact of non-autonomous LTR retrotransposons

**DOI:** 10.64898/2026.03.01.708851

**Authors:** Sophie Maiwald, Ferdinand Maiwald, Tony Heitkam

**Author notes:** corresponding authors: Tony Heitkam, Sophie Maiwald.

## Abstract

Plant genomes are filled with retrotransposons and their derivatives, subject to constant sequence turnover. As short, non-autonomous retrotransposons do not encode a protein product, they experience reduced selective constraints on their DNA sequence, leading to diversification into multiple families, usually limited to only a few species. This absence of any coding capacity and their tendency to form subfamilies are the reasons for the incomplete description of non-autonomous LTR retrotransposons in most to all genomic repeat annotations. Here, we focus on non-autonomous LTR retrotransposon identification. Are all of these sequences derivatives of easier-to-identify full-length elements? Or is there more variability, which is currently overlooked? For this, we capitalize on our comprehensive understanding of the TE landscape in sugar beet to assess the extent of the blind spot on non-autonomous LTR retrotransposons

Here, we present a workflow to identify non-autonomous LTR retrotransposons without prior sequence information, retrieving more than 100 families within the sugar beet genome. We only include TEs without the ability for complete self mobilization. Spanning up to 15,000 bp, these non-autonomous families are often longer than expected and characterized by reshuffling and modular evolution. Most strikingly, only a few of these families are directly derived from autonomous partners, showing that there is a large, undiscovered TE variety in the non-autonomous TE fraction.

We highlight that a large fraction of non-autonomous TEs won’t be retrieved with the current TE identification workflows, even if the output is well-curated and condensed into TE libraries and suggest procedures to remedy this gap. This study is the first insight into the non-autonomous LTR retrotransposon landscape within a single genome and serves as an example to estimate the error in non-autonomous TE detection.

## Introduction

Large fractions of plant genomes are formed by repetitive DNA units of all kinds. Mobile genetic elements, or transposable elements (TEs), accumulate in the genome [1–3] and have a major impact on genomic infrastructure and integrity [4,5]. Due to their ability to actively change their position and therefore occupy multiple genomic locations, they are a source of genomic reorganization [6–8]. Based on their structural hallmarks, order of mobility-allowing protein domains, and the resulting movement processes, transposable elements are classified into multiple groups.

Here, we focus on long terminal repeat retrotransposons, a large group of transposable elements (TEs) propagating via a copy-and-paste mechanism with the help of an RNA intermediate [9]. Despite their great variety, the overall structure is highly conserved and delimited by two identical flanking regions (a result of the copy-and-paste mechanism), the *long terminal repeats* (LTRs). LTRs often contain signals for transcription initiation and termination [10–12]. The internal region between the LTRs contain the conserved binding sites for reverse transcription (*primer binding site* - PBS and *polypurine tract* - PPT) and conserved protein domains for reverse transcription and integration (RT - *reverse transcriptase*, INT - *Integrase*, PROT - *protease*, RH - *ribonuclease H* and GAG). Usually, LTR retrotransposons are condensed into families based on the order and similarity of the coding domains [13].

Nevertheless, with a closer look, a great variety of short LTR retrotransposons without any reminiscence to the coding regions can be identified. Similar to their autonomous counterparts, non-autonomous LTR retrotransposons are characterized by identical flanking LTRs as well as PBS and PPT regions. Traditionally, these non-autonomous LTR retrotransposons have been subdivided based on their overall sequences length: whereas *terminal retrotransposons in miniature* (TRIMs) [14] are described as short (< 1 kb), *large retrotransposon derivatives* (LARDs) [15] usually are long (> 4 kb). For the purpose of manuscript, we will use TRIM as a unified designation for all non-autonomous, non-coding LTR retrotransposons, regardless of their length.

Due to their lack of a coding region, TRIMs are typically shorter than their autonomous counterparts. Hence, they may be more easily tolerated in the genome, potentially also in more actively transcribed regions. Similarly, their short length favors recombination, solo-LTR and tandem formation, and thus may contribute to an overall increased structural variation [15–18].

TRIMs are difficult to identify in genome sequences. This is because their rather low complexity makes them unsuitable targets for the current TE identification and classification approaches, which are typically either similarity- or signature- (structure-) based or a combination of both [19–21]. We identify the following three hurdles in TRIM identification:

1. Similarity-dependent TE tools fail: These tools search for known TE protein domains in genome assemblies. This is possible as these coding regions show high conservation in nucleotide/amino acid composition across different species [22–24]. For TRIMs, as these conserved protein domains are missing, the chance of identification is low and only possible if their LTRs show at least partial similarity to already described autonomous LTR regions.
2. *De novo* identification produces many false-positives: As similarity-based methods fail, only a *de novo* identification is possible, combined with signature-based approaches. Signature-based identification scans the genome for coding domains and smaller conserved motifs, such as PBS sequences, combined with an all-by-all sequence comparison to identify repeated regions such as LTR sequences. Several signature-based tools for LTR *de novo* identification are available [25–27] with LTR_Finder [28] probably being the most known and best-performing (although being slow) as described by Ou et al., 2019 [22]. The advantage of these signature-based tools is their ability to identify LTR retrotransposons not only based on the presence of coding or conserved motifs, but by the presence of identical flanking regions in the form of LTR sequences. Hence, this approach is suitable for *de novo* identification of non-autonomous LTR retroelements. On the downside, this process still needs a lot of manual curation as the availability of LTR sequences as the only structural and describing hallmark of these elements leads to a high amount of false-positive hits, especially as truncated or tandemly-arranged copies are also detected [29]. Especially in the case of non-autonomous LTR retrotransposons, manual curation is often waived and as a result, these retrotransposons are mostly not included in TE databases.
3. Manual curation is impeded due to vague TRIM boundaries: The correct annotation of full length retrotransposons depends on the correct assessment of the TE’s boundaries. This is especially difficult for non-autonomous elements. Self-comparison of sequences is an efficient strategy for boundary detection. First introduced in 1970 by Gibbs and McIntyre [30], dotplots are an efficient way for self-comparing sequence information allowing detection and annotation of various, even complex repeat types and modular similarity [20,31–33]. Furthermore, dotplots describe sequence information in a way which tackles the fine line of being human- and machine-readable at the same time. This allows manual curation as well as further image processing with one output.

In summary, despite their potential to contribute to genome regulation and structural variation, we still have barely investigated non-autonomous LTR retrotransposons. Their rather limited sequence complexity is at fault, leading to an under-annotation in genome sequences. The TE community is largely aware that this annotation gap exists, but it is unclear if this gap is large or negligible. We here aim to evaluate the size of the blind spot that we introduce into our studies, if we disregard TRIMs. For this, we will fully identify and characterize TRIMs in a moderately-sized plant genome, check which fraction is typically missed and evaluate how this may impact studies of genome regulation and structural variation. As a genomic reference for our TRIM study, we chose the genome sequence of sugar beet (*B. vulgaris*) as its TE landscape is very well-understood including repetitive DNA elements of both TE classes (summarized in [8,34]), detailed in-depth studies of different LTR retrotransposons [35–37] and non-autonomous elements [18,38]. It is a moderately-sized (758 Mb) genome with a high-quality long read-based genome assembly [39].

Here, we present a workflow for the *de novo* TE identification for non-autonomous LTR retrotransposons with a focus on identification and quantification. We discuss in detail which tools and parameters are useful and needed to get an impression of the non-autonomous LTR retrotransposon landscape. Furthermore, we elaborate on classification of these elements and the obstacles one has to tackle when working with non-coding repetitive elements. Our comprehensive study focusing solely on non-autonomous LTR retrotransposons enables a deeper look into the broad variety of these often neglected and vastly underestimated TEs and opens the way for a deeper understanding of genome dynamics in the light of non-autonomous LTR retrotransposon evolution.

## Material and Methods

### *de novo* non-autonomous LTR retrotransposon identification

For first identification of non-autonomous LTR retrotransposons we performed an LTR_Finder analysis. As non-autonomous LTR retroelements do not encode any protein domains, we chose LTR_Finder settings avoiding any coding regions as a necessary condition. Hence, we also avoided the conserved 5’-TG..CA-3’ di-nucleotides framing the LTR sequences. Therefore, motif search of LTR_Finder was limited to only the occurrence of PBS and PPT (-F 0000011000). For PBS prediction we were using motifs from the plant tRNA database (http://seve.ibmp.unistra.fr/plantrna/ [40]). As we were interested in the whole landscape of non-autonomous elements in sugar beet, we tested three different parameter settings. We knew from previous test runs, default LTR_Finder settings are not suitable for *de novo* identification of very short sequences. As input, we used an Oxford Nanopore Technologies based genome assembly of sugar beet, *B. vulgaris* (GCA_947631125 [39]).

We decided to test three stringency parameters in the following ways (Table 1):

1. “Run1 - stringent”: We targeted short non-autonomous sequences with the strict parameter setting suggested by Gao et al., 2016 [17] that assumed very short LTR sequences and short overall TE length.
2. “Run2 - less stringent”: To identify non-autonomous LTR-retrotransposons of all lengths, we decided to relax stringency regarding LTR distance.
3. “Run3 - relaxed”: We further relaxed the parameters for maximal LTR length and distance. As different LTR_Finder parameters lead to the detection of identical families, we used an intersection of all runs for further analysis and quantification to reduce duplicated reference sequences.

**Table 1:** Parameter settings used for LTR_Finder analysis. LTR_Finder runs are sorted by stringency for short non-autonomous LTR-retrotransposons.

| Run | stringent | less stringent | relaxed |
| --- | --- | --- | --- |
| min. LTR distance [ bp] | 30 | 30 | 30 |
| max. LTR distance [ bp] | 2,000 | 6,000 | 10,000 |
| min. LTR length [ bp] | 30 | 30 | 30 |
| max LTR lengths [ bp] | 500 | 2,500 | 5,000 |

### Output processing and first tier filtering

The LTR_Finder output was processed into *.gff* file format with Python3. The converted LTR annotations were inspected in Geneious Prime (https://www.geneious.com) and the resembling nucleotide information was extracted. For output filtering we performed a BLASTx analysis against described protein coding domains from the REXdb Viridiplantae v3.0 database (http://repeatexplorer.org/?page_id=918 [12]).

### Second tier filtering: Graphical dotplot filtering and annotation of suitable candidates

To reduce false positive hits, we assessed the structural intactness of the TRIM candidates by introducing a graphical filtering step: Filtered LTR_Finder outputs were plotted as self dotplots with Flexidot v1.06 [41]. Because of the shorter length and high variability in the dataset, smaller word sizes are more suitable when plotting these sequences. Therefore, we adjusted the flexidot plotting outputs with -f 1 -k 7 -p 0 -r N -E 7. For faster processing of all dotplots we used a computer vision-based approach to search for LTR retrotransposon-specific dotplot patterns, in detail laid out in Additional Fig. 1. Shortly, flexidot outputs are split and binarized using Otsu thresholding [42]. In the next step contours are detected and filtered by size. For noise reduction we use morphological operations. For each plot the largest contours are analyzed based on geometric properties (e.g. width, area, position) to distinguish different types of structural arrangement. Sequences with no significant contours are filtered out.

**Fig. 1:**
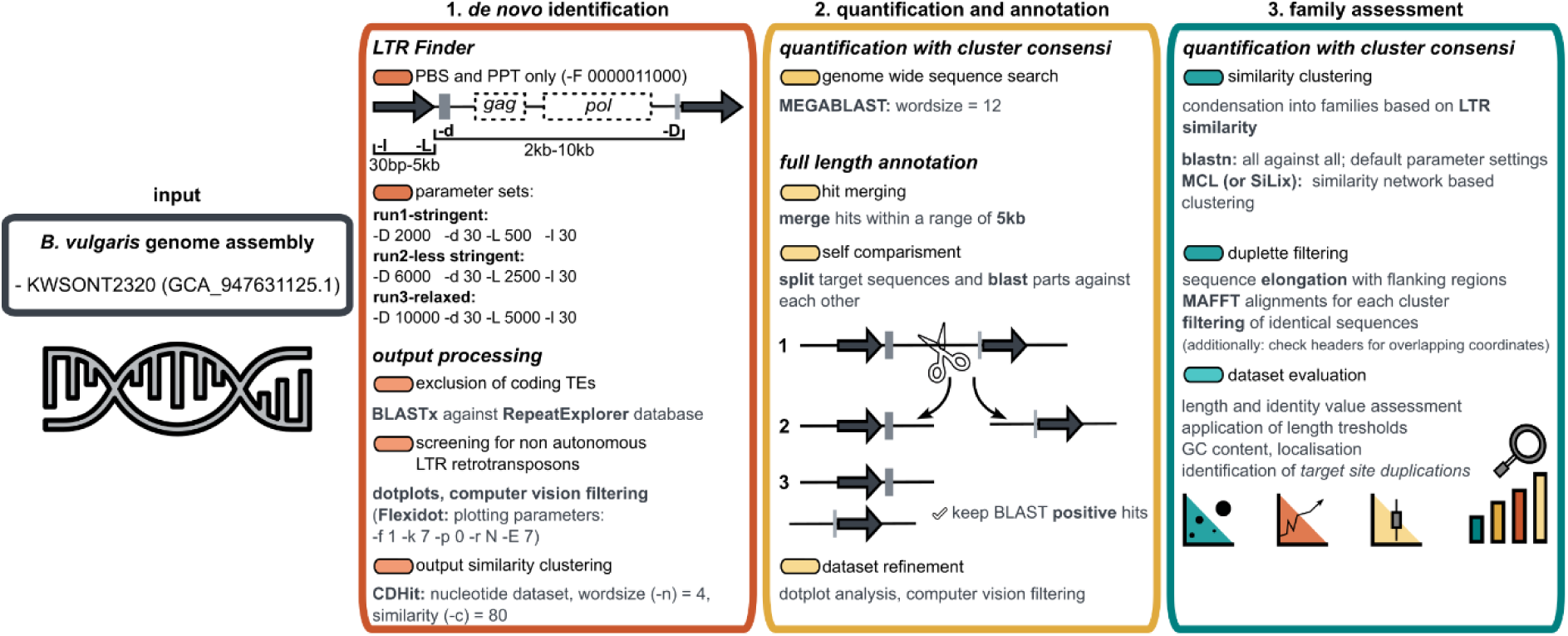
Three steps to identify non-autonomous LTR retrotransposons. As *input* we used an oxford nanopore technologies (ONT)-based genome assembly of *Beta vulgaris*. Assessment of non-autonomous full length sequences was performed with three main steps: *Step 1* - *de novo* identification and output processing; *Step 2* - quantification and annotation of full length sequences; *Step 3* - family assessment via clustering and metadata evaluation.

The evaluated reference LTR retroelements then were inspected in Geneious. This TE annotation step entails identification of all internal domains, such as LTR, PBS and PPT. Additionally, reference sequences were clustered using CDhit [43] for nucleotide sequences and similarity values adapted to less conserved TEs (-c 0.8). Reference sequences (either single or representatives for each cluster) were used for quantification.

### Quantification and annotation of full length sequences

To reduce noise and yield highly similar hits we performed quantification via local Megablast [44] with reference sequences as a custom library on the KWS2320ONT assembly and a wordsize of 12. MEGABLAST calculation was done with the open source bioinformatics platform galaxy (www.usegalaxy.org). The following steps of file processing and hit extraction were performed with custom python and R scripts. The quantification output was converted to .gff file format and hits within a range of 5 kb were merged to enable assessment of tandemly arranged copies. Extraction of fasta sequences was performed with and without a range of flanking regions for identification of target site duplications (TSDs). Dataset reduction was performed with filtering against coding domains as described above. To reduce the dataset even more and get rid of solo and single LTRs or highly truncated copies, we added an additional step of sequence self comparison for each single sequence. For this, extracted hits were cut on a calculated middle position, generating left/right pairs for each sequence. Then left and right parts were blasted against each other and all hits without positive self comparing signals were excluded from the dataset. Hits larger than 10 kb were cut into quarters, as we suspected a tandem arrangement of these sequences due to their size. Final annotation of full length sequences was performed as described above with a dotplot based approach and manual curation.

### Clustering and family assignment

To understand the biology and genomic behavior of non-autonomous elements, classification in form of similarity-driven condensation into families is beneficial. Classification of LTR retrotransposons is mostly performed on autonomous elements with coding regions. Hence, we tried applying several approaches on our dataset relying on LTR sequence nucleotide similarity to check which performs best for non-autonomous elements.

#### 1.) kmer based identity evaluation

K-mer calculated similarities are often used and well established within the research community. In our tests we used two different tools - CD-Hit [45] and MMseqs2 [46] and compared their performance. Both provide fast and efficient dataset clustering with different strategies: CD-Hit performs pairwise k-mer similarity assessment followed by a greedy clustering approach, whereas MMseqs2 uses adaptive k-mer filtering and indexing to approximate sequence similarity. We tested multiple parameter sets for each tool: (1) k-mer word size (CD-HIT: n=4–9), (2) minimum sequence identity (CD-HIT: i = 80–90%; MMseqs2: i = 70–90%), and (3) minimum coverage/length overlap (CD-HIT: l = 70–90%; MMseqs2: l = 70–90%).

#### 2.) Similarity-network based clustering

Due to observed LTR sequence appearance, broad length varieties and alignment-based (e.g. with MUSCLE or MAFFT) similarity calculations are unsuitable. Instead, we performed an all against all BLAST (blastn; default parameters) and used the results for similarity-network clustering. We tested two similarity graph based clustering algorithms: SiLix (Single Linkage Clustering of sequences [47]) uses a single connectivity within the graph under coverage constraint to build clusters, whereas MCL (Markov Clustering on similarity graphs [48] is clustering sequences by simulated random walks normalized with a Markov matrix for transition probabilities. These are controlled by the inflation parameter, which corroborates strong connections within clusters and weakens inter-cluster noise. We chose various pairwise similarity and alignment coverage thresholds for SiLix and MMseqs2 all against all search (identity: i = 70-90%, coverage: l = 70-90%). For MCL we explored a range of inflation parameters (inflation: in = 2-5).

Cluster evaluation was performed within Geneious and manual curation. Due to modular organisation of non-autonomous LTR retrotransposons similarities occur across family borders leading to duplicated hits. Duplicated hits were excluded from the dataset based on their coordinates on the pseudochromosomes of the sugar beet assembly or nucleotide composition including flanking regions. As LTR-retrotransposons populations expand via a copy and paste mechanism the occurrence of identical copies on multiple genomic loci is possible. Comparing full length sequences and their flanking regions enables identification of true positive duplicated hits within each cluster.

##### Target site duplication (TSD) identification

Identical motifs in form of *target site duplications* (TSD), which are a result of active transposition can be located upstream and downstream of the element borders with a length of 5 nt. For automated identification of TSDs, we used a custom Python routine, which performs based on existing annotations resulting from full length filtering. We allow 1 mismatch, defining TSDs as 5 bp long sequence motifs with at least 80% identity.

##### Annotation of tandemly-arranged sequences and single LTRs

Results of recombination such as tandemly-arranged sequences and single LTRs were identified with different strategies. For the first, we scanned dotplots for patterns indicating a tandem arrangement. The underlying sequences were then blasted against reference sequences from each family to determine family-specific assignment. For single LTR annotation, we used the “solo_finder.pl” plot from the LTR-retriever pipeline [49]. For both outputs, the final annotation of single LTRs was performed manually.

##### Filtering of truncated or re-arranged copies

To identify structurally altered TRIM-derived sequences beyond canonical full length elements, solo-LTRs and tandemly-arranged copies, we used BLAST-based filtering and summarization. First, all full length copies as well as solo-LTRs were excluded from the initial MEGABLAST quantification dataset. Remaining filtered sequences, representing putative truncated and rearranged derivatives, were then mapped locally against a curated TRIM reference dataset using MEGABLAST (BLAST+ blastn, task megablast). BLAST search was performed with an increased maximum target sequence limit of 20,000 to avoid truncation of high-copy families.

BLAST hits were grouped by reference query sequence, and summary statistics were calculated for each non-autonomous family. These included the total number of alignment hits, the number of unique subject identifiers, and the number of unique genomic loci, the latter determined by parsing contig identifiers and genomic coordinates embedded in subject sequence headers. For each family, mean percent identity, mean alignment length, mean bitscore, and the highest-scoring alignment were also recorded. This approach allowed quantification of partially conserved TRIM-derived sequences that retain detectable similarity to reference elements but lack the structural features required for classification as intact TRIMs or solo-LTRs, providing a genome-wide measure of recombination-driven truncation and rearrangement.

##### Gene association study

Localization of TRIM sequences as well as autonomous LTR retrotransposons from sugar beet [34] were illustrated with the circos package within R circlize [50]. Additionally, we checked for gene association of our TRIM dataset with a published gene annotation of a reference sugar beet genome [39] with bedtools [51] for direct overlapping (bedtools intersect - a <TRIM dataset> -b <GENE dataset> -wa -wb > <INTERSECT output file>) as well as within a certain window of 5 kb to check association in gene flanking regions (extend genes by 5 kb each direction: bedtools slop -i <GENE dataset> -g genome.sizes -l 5000 -r 5000 > <FLANK output file>; remove gene bodies: bedtools subtract -a <FLANK output file> -b <GENE dataset> > <ONLY flank output file>; intersect: bedtools intersect -a <ONLY flank output file> -b <TRIM dataset> > <INTERSECT with flank output>).

## Results

### Detection of non-autonomous elements on a genome-wide level

To investigate non-autonomous LTR retrotransposons two major questions arise:

1.) “How many non-autonomous LTR retrotransposon families reside in a typical plant genome?”; and 2.) “How effectively are these elements detected by current pipelines for repeat identification?”

To provide a basis for both questions, we aim to fully identify all non-autonomous LTR retrotransposons (= terminal-inverted repeat retrotransposons in miniature, TRIMs) in a well-studied, moderately sized plant genome. For this, we chose the sugar beet genome as we have a deep understanding of its LTR retrotransposon landscape [18,34–37,52]. Our procedure is structured into three major steps as shown in Fig. 1: (1) *de novo* identification, (2) quantification and annotation and (3) family assessment.

Starting with the first step, the core of non-autonomous LTR retrotransposon identification is the prediction of repeats based on structural features. So-called signature based approaches rely on the availability of conserved motifs, in the case of non-autonomous elements tRNA-derived *primer binding sites* and structural features like *long terminal repeats* (LTRs) and *target site duplications* (TSDs). Our tool of choice was LTR_Finder, which is well established within the repeat community.

To capture the whole length spectrum, we decided on three different parameter sets: (1) The parameter combination “specific and stringent” for short non-autonomous LTR retrotransposons as described in previous studies [17,18], (2) a less stringent set with expanded length ranges approximating LTR_Finder default settings, and (3) a relaxed configuration to detect very large elements. For all three runs, it is crucial to disable LTR_Finder to detect LTR retrotransposon specific protein domains, as non-autonomous elements lack these.

As expected, LTR_Finder yielded hits with low scores and hit length distributions resembling the chosen length parameters (Additional Fig. 2). Notably, for non-autonomous elements, aiming for low scores is beneficial, as the score reflects the presence of coding motifs and protein domains. Initial hit numbers ranged from 6,000 (stringent) up to 39,000 (relaxed) and were subsequently reduced through filtering and annotation steps.

**Fig. 2:**
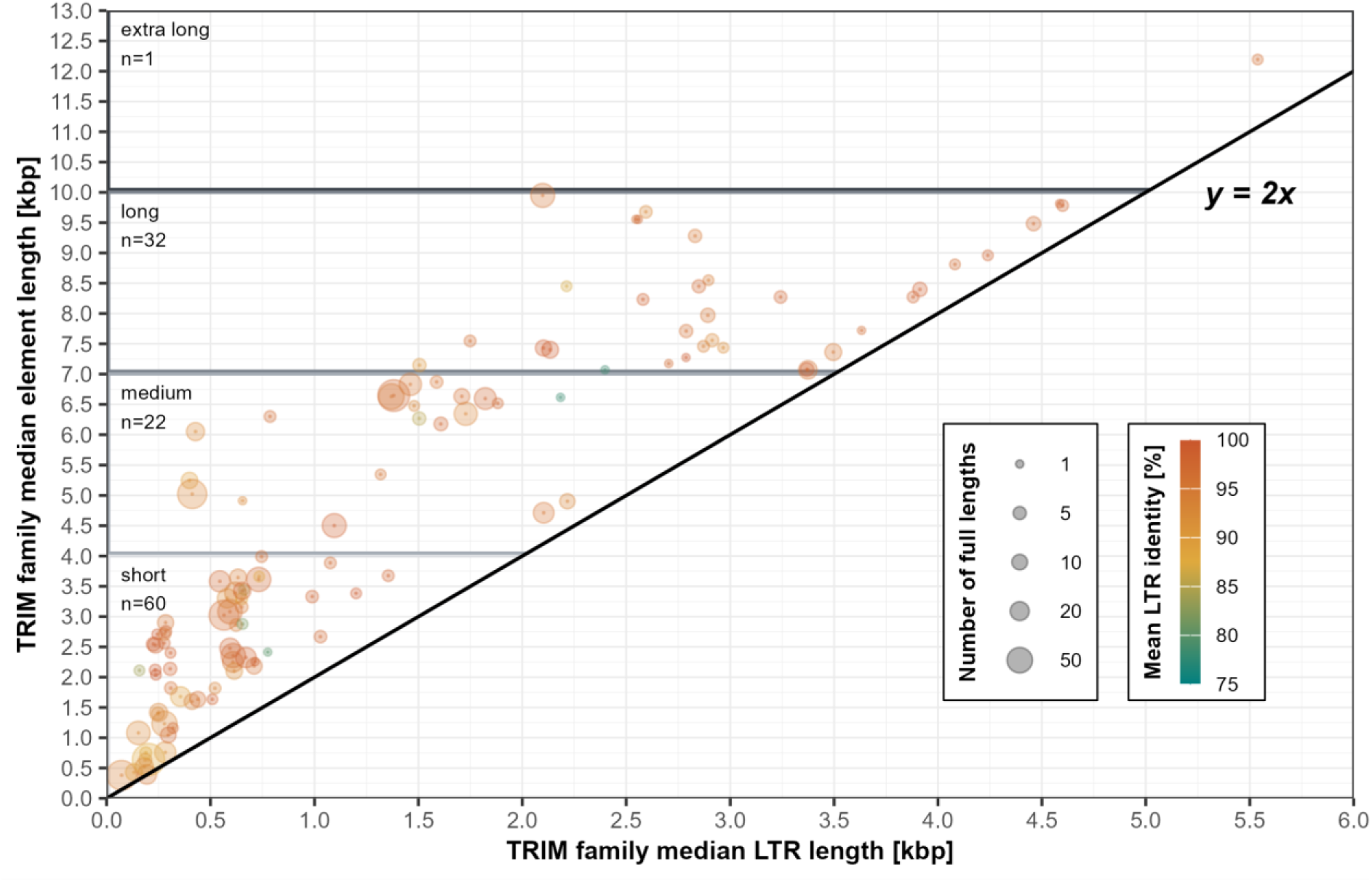
Length distributions across non-autonomous elements in sugar beet. Shown are all calculated median LTR and corresponding full length values for each TRIM family. Dots are scaled according to the number of full length sequences representing each family. For every sequence within the dataset, the pairwise LTR identity was calculated and their mean was included for each family: blue coloring indicating low values and orange colouring corresponding to high mean LTR identities. The blackline refers to LTR and full length combinations, arithmetically impossible to reach as they are smaller than the doubled LTR size (y = 2x).

As LTR_Finder still detected some hits with coding regions in the less stringent and relaxed runs, we excluded these by filtering with a BLAST against the coding domain database of known LTR retrotransposons available via the RepeatExplorer platform. To get an impression of sequence structures, we calculated dotplots for all of them and scanned them with a computer vision based approach. This was beneficial for two reasons: 1.) dotplots provide visual overview on structural arrangement of a nucleotide array which is both human- **and** machine-readable. 2.) Automated scanning for LTR retrotransposon-specific patterns accelerates manual sorting and curation. Supported by dotplots, we manually curated the dataset and eliminated false-positives. Further artifacts such as truncated, deleted or otherwise mutated hits were eliminated after application of a clustering step. We used CDHit for quick and efficient initial clustering with a wordsize of 4 and 80% nucleotide similarity for each cluster per run. Ultimately, we used 252 reference sequences for quantification. These were derived from an intersection of all non-autonomous LTR retrotransposon-positive cluster consensi across the stringent (n=136), less stringent (n=322) and relaxed (n=333) LTR_Finder runs.

For quantification (Fig. 1, step 2), we decided on MEGABLAST with a reduced wordsize of 12. This adjustment increases sensitivity, making it suitable for detecting shorter non-autonomous LTR retrotransposon families and high-throughput screening while still being computationally efficient and alignment-sensitive. As the primary goal of our study was to get a more comprehensive view of the general organization and appearance of non-autonomous LTR retrotransposon landscapes, we initially focused on full length sequence annotation. Beforehand, we recommend an additional filter step against protein-coding LTR retrotransposon domains to avoid detection of autonomous elements similar to the targeted non-autonomous ones. Noise such as solo-LTRs, fragmented artifacts and truncated elements was reduced with a self-BLASTing filtering step. The model for this approach was the detection of intact elements done by the DANTE-LTR pipeline [24], in which LTRs are detected by aligning flanking regions up- and downstream of identified coding protein-domains. As our focus are non-autonomous elements, we adapted this strategy by splitting each MEGABLAST output hit at the midpoint into pairs of left and right. Self-blasting of left/right pairs led to a reduction of candidate sequences from 350,997 to 31,476. For detailed full length annotation, we returned to the procedure described above: generating dotplots, assessing them based on defined structural features, and performing manual curation of primer binding site and polypurine tract. Furthermore, sequences were scanned for intact LTR sequences (5’-TG…CA-3’) and large indels within the LTRs to guarantee the presence of only complete non-autonomous LTR retrotransposons in the dataset.

In total, 7,182 intact non-autonomous full length LTR retrotransposons needed condensation into families. For this, clustering was performed based on LTR sequences, as these are the most conserved family-specific features detectable across all sequences. After manual curation, we identified 115 families, which served as ground truth for evaluation of clustering tools and parameter tests. Detailed information for cluster numbers of tested parameters is compiled in Additional Fig. 3 and summarized as follows: All-against-all BLAST followed by Markov Clustering Algorithm (MCL) yielded cluster numbers closest to the ground truth, across both relaxed and stringent parameters. Manual inspection confirmed that clusters were consistent and family-specific. In contrast, we observe strong overclustering for the approaches combining all-against-all BLAST and SiLix as well as MMSeqs2 all-against-all search followed by MCL. These methods produced more clusters than the ground truth, often splitting true families into multiple smaller clusters or singletons. All k-mer based approaches and parameter combinations resulted in severe overclustering, producing high cluster numbers and a high amount of single sequence clusters. These outcomes are a result of excessive fragmentation of true families.

**Fig. 3:**
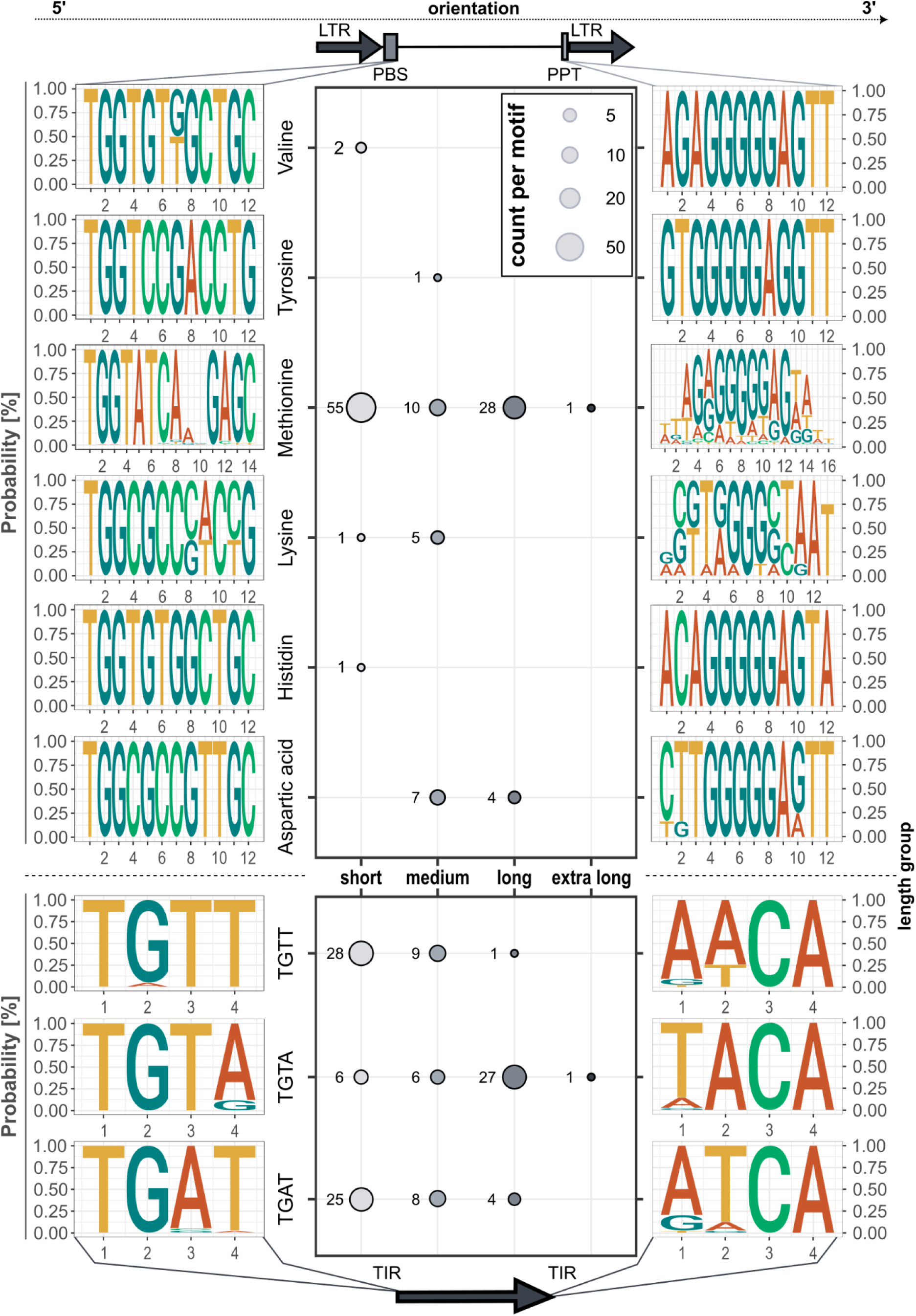
Analysis of identified primer binding site (PBS) motifs and the corresponding tRNA, polypurine tracts (PPT) and terminal inverted repeats (TIR). All identified tRNA-derived and TIR motifs and their corresponding 3’ PPT motifs as well as terminal inverted repeat (TIR) motifs are shown as sequence logos. The amount of families with a certain motif are listed regarding their length groups.

The final dataset of non-autonomous LTR retrotransposons after annotation and applying strict length thresholds comprises 1,581 full length sequences in 115 families with up to 95 full length sequences within.

Addressing our initial questions on recent repeat annotation tools and how they cope with non-autonomous LTR retrotransposons detection: Out of our 115 detected non-autonomous families in the final dataset 86 were recognized by LTR_Finder with default settings (enabling detection of protein coding domains). A better result was yielded for the recent *de novo* identification pipeline EDTA, which harvested 97 families. Unfortunately, all of the identified TRIM families were identified as LTR/unknown by EDTA which leads to a combined dataset of TRIMs together with recombined, not intact or otherwise rearranged LTR retrotransposons, resulting again in manual curation.

### The TRIM population is highly diverse: 42-fold differences in length and varying motifs

The first aspect within this newly defined non-autonomous dataset was length and size distributions of TRIMs. We observe a considerable variety of overall lengths (Fig. 2). TRIM-01 represents the shortest family, with a median overall length of 291 bp and median LTR length of 72 bp. At the opposite end of the spectrum, TRIM-127 is the largest family, with a median length of 12,284 nt and median LTRs measuring 5,533 bp. Across the dataset, family-specific median lengths and LTR sizes span this entire range, although the majority of families exhibit a median overall length of less than 4 kb. For data evaluation purposes, we decided to divide the dataset based on overall median length and defined four groups (Fig. 2): short (0 - 4 kb), medium (>4 kb - <7 kb), long (>7 kb - <10 kb) and extra long (>10 kb).

No consistent relationship was observed between overall element size and related lengths of the LTRs and internal regions. Larger elements do not necessarily correspond to longer LTRs. For example, TRIM-06 with an overall length of 5.5 kb and LTR lengths of 186 bp, resulting in a length/LTR ratio of almost 15.

Length/LTR ratios higher than 3.5 indicating comparatively long internal regions, were detected in 17 families. In 10 (TRIM-03,04,11,12,14,16,19,20,33,34) the elevated ratios can be attributed to repeat arrays embedded in the internal region. These arrays occur in a repeated, satellite-like configuration (Additional figure 4). In contrast, 65 families exhibited small ratios of 2 or less, reflecting relatively short internal regions. Although no universal scaling rule emerges, a general tendency is visible: shorter internal regions are more common among families with greater overall length. This is particularly evident in families with LTR lengths of at least 2.1 kb, where ratios fall to 1.75 and less (Additional Table 1.1).

**Fig. 4:**
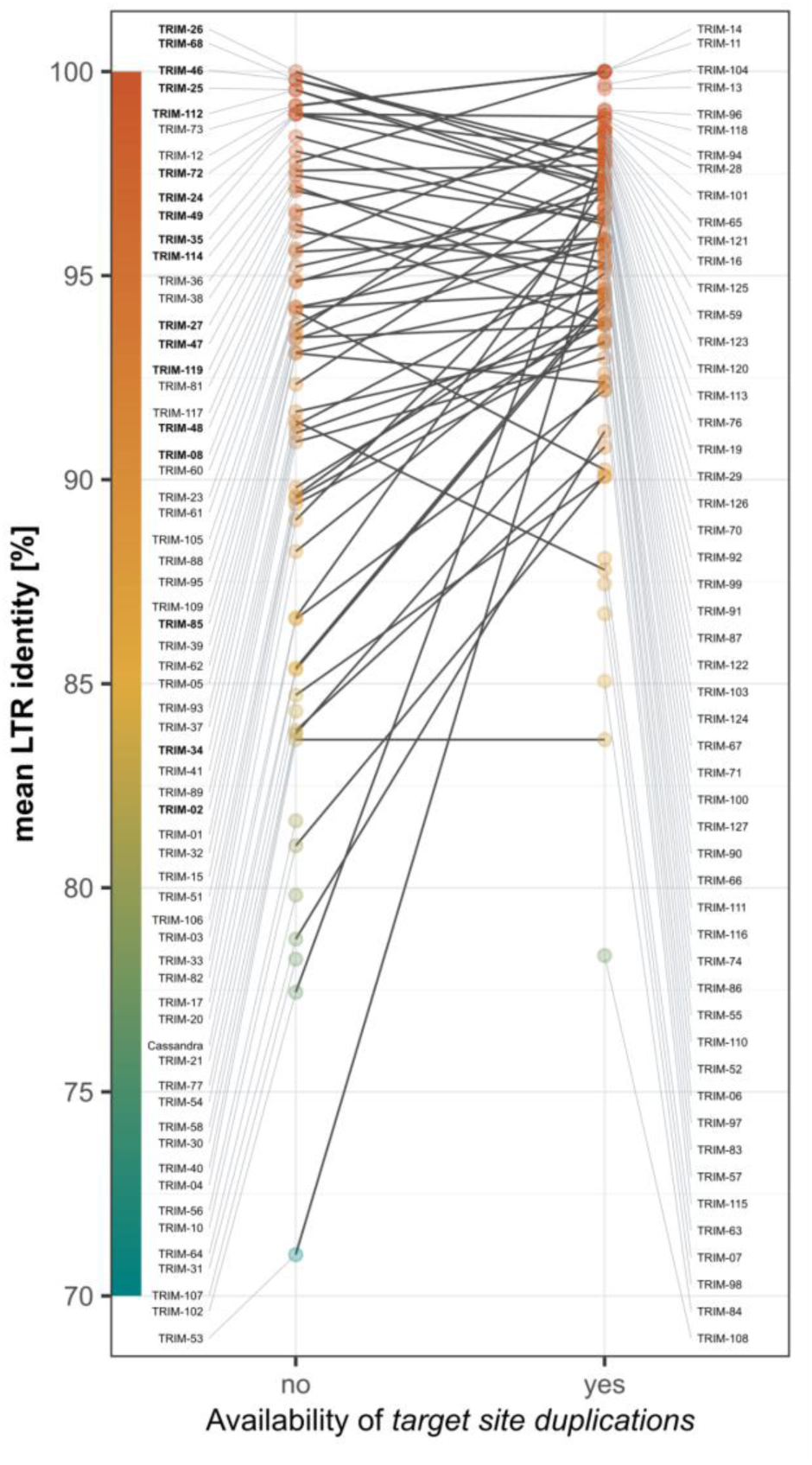
Comparison of *target site duplication* (TSD) availability and mean LTR identities per family. Each dot represents a TRIM family fractions with and without TSD based on their mean LTR identity. Higher identity values are shaded in orange. Labelled in bold are families with higher mean identity values for full length copies without TSDs compared to those with.

Interestingly, we observe high median pairwise LTR comparison values across the whole population. The majority of families show high similarity with LTR identity values of 85% and higher (Fig. 2, color gradient). Hence, only five very small families with observed full length sequence numbers of 2 or less represent lower median LTR identities (TRIM-04, 56, 58, 64, 107).

Although non-coding, non-autonomous elements retain conserved motifs required for reverse transcription, such as the *primer binding site* (PBS). We analyzed these motifs and assigned the corresponding tRNAs. Together with the *terminal inverted repeats* (TIRs) defining LTR boundaries, these features can be valuable indicators for superfamilies from which these non-autonomous elements originated. In total, we identified six different PBS sequence motifs across all 115 families (Fig. 3). Among those, one is derived from histidine tRNAs, one is derived from tyrosine tRNAs and two from valine tRNAs. Lysine and aspartic acid derived PBS sequences are present in 6 and 11 families respectively, predominantly present in medium-sized TRIM families. Methionine-derived PBS sequences are the most common and distributed across the whole TRIM length spectrum.

We also assessed the corresponding polypurine tracts (PPTs), but no clear signals or corresponding motifs are detectable (Fig. 3). Regarding TIR sequences, we identified three different 4nt long motifs forming LTR borders, which were identified across the defined length groups. For instance, 5’-TGTT-3’ and 5’-TGAT-3’ are most common in short families, while 5’-TGTA-3’ is dominant, but not exclusively present in longer families.

### Target site duplications indicate recent integration

Age estimation for LTR retrotransposons can be a challenging task as it is dependent on substitution rates that are usually deduced from coding or highly conserved genic regions, which evolve differently from mobile DNA elements e.g. in the light of repeat homogenization [53]. Furthermore, substitution rates are often species-specific [54]. As an alternative, we used pairwise calculated LTR identities as an indicator for conservation and dynamics of retrotransposon families. In general, higher LTR identities indicate more recent insertions, as the two LTR sequences are still nearly identical shortly after integration.

In our dataset, 79 families show mean LTR identities of 95% or higher, suggesting recent insertions. On the other hand, only two families exhibit mean LTR identities of 80% and lower, consistent with the assumption of older populations accumulating mutations within the LTRs and becoming unrecognizable in the genome. Another hallmark for successful (and typically more recent) insertion is the presence of *target site duplications* (TSDs).

Across all 1,581 full-length TRIM sequences, only 229 lack a detectable TSD motif. Five families only contain members entirely without TSD sequences; these families are small, comprising only seven full-length sequences in total. Elements from families with lower LTR identities (< 85%) most often lack TSDs, supporting the idea that older elements accumulate mutations in both their LTRs but also the flanking regions leading to the erosion of recognizable TSDs over time.

This contrasts with the finding that non-autonomous full length copies from families (n=11) with LTR similarities of >=95% lack TSD motifs, likely an indicator of genomic rearrangements and recombination. This shows that the presence of TSDs is not universal across TRIMs, even among the most recent insertions. Overall, 108 out of 115 most of the families contain at least some TSD-carrying sequences, highlighting their general prevalence among TRIMs.

### Recombination is common in non-autonomous retrotransposon biology

To explore the impact of recombination on the TRIM landscape, we analyzed the abundance of TRIM rearrangements, especially tandemly arranged (TA) TRIM copies and single LTRs (solo-LTRs) as well as modular relationships between different TRIM families.

We observed tandemly repeated TRIMs, both with and without associated TSDs. In total, we identified 97 TA copies across 38 families, with the number of LTRs per tandem ranging from 3 up to 21. The major fraction of TA copies (65) displays a 3-LTR-structure. 15 out of the 38 families contain only a single TA copy, while the remaining show multiple tandem formations. 73 out of the 97 identified TA copies retain identifiable TSDs. There is no clear relationship between tandem formation and total copy number or high LTR similarity: tandem formation was observed in both high-copy families (e.g. TRIM-01) and single-copy families (TRIM-65, TRIM-125) and families with LTR identities below 90%.

Evidence for recombination is also apparent from the presence of single LTRs; again, either with or without flanking TSD sequences. For solo-LTR identification we used two independent methods: 1.) by filtering the MEGABLAST output used for full length quantification based on hit position and alignment range and 2.) by using the integrated solo-LTR identification module from the LTR_retriever pipeline. Both approaches yielded consistent results. In total, 101 of 115 families show evidence of solo-LTRs, while 14 families lack them entirely. Solo-LTR abundance varies widely, with “solo-LTR : full length” - ratios ranging from 0.01 up to 111. For instance, TRIM-01 is one of the most abundant families with 77 full length copies but shows the lowest ratio of 0.01 with only one solo-LTR, whereas TRIM-26 exhibits 223 solo-LTRs but only two canonical full length copies, resulting in the highest ratio of 111.5. Among the families with solo-LTRs, 54 have at least 50% of their solo-LTRs flanked by TSDs. Of these, 20 families also contain TA copies and 18 show high TSD retention of at least 50%.

As described in the methods section, we applied a length threshold of +/- 15% median length to all non-autonomous families to define full length sequences. For some families, length deprivation between single members is volatile due to formation of subpopulation, e.g. for TRIM-116 length variation even exceeding 15% (Additional Tab. 2) because modular similarity across the whole family leads to manual condensation into one family.

Additionally, we observe shared sequence similarity between TRIM families. Families with shared similarities were condensed into “modular groups” (1-16) and analyzed regarding shared regions (Additional Fig. 5). For modular groups M6 and M7 we additionally identified one TA copy, each not clearly assignable to either one of the corresponding families. Such chimeric elements are not limited to families with shared similarity, e.g. two chimeric elements are assignable to TRIM-08 with additional LTRs donated by either TRIM-01 or TRIM-05 copies.

In addition to canonical full-length elements, tandem arrays, and solo-LTRs, we detected widespread evidence for structurally altered TRIM-derived sequences that retain partial similarity to curated reference elements. To quantify these derivatives, we mapped the filtered MEGABLAST hit output, after exclusion of full-length TRIMs and solo-LTR candidates, against the TRIM reference sequences and summarized alignment statistics per reference (Additional Table 1.2). For many families, large numbers of hits remain detectable despite failing to meet structural criteria for intact elements, consistent with truncated copies, or copies with insertions/deletions.

The abundance of such altered derivatives varies among non-autonomous families. For example, TRIM-01, one of the most abundant families in the genome (full length-wise), shows more than 1,600 unique hits with detectable similarity to the reference sequence. Similarly, TRIM-33, TRIM-49, TRIM-65, and TRIM-116 each exhibit several thousand non-canonical hits, indicating extensive accumulation of fragmented or rearranged copies. In contrast, low-copy families such as TRIM-06, TRIM-07, or TRIM-14 show only a few dozen hits.

Despite their structural heterogeneity, sequence similarity across remaining hits remains relatively high, with mean nucleotide identities frequently ranging between 85% and 95%, indicating that these derivatives are trusted assignments to the corresponding reference sequences.

The prevalence of truncated or rearranged copies is not restricted to families with high abundance of tandemly-arranged copies or solo-LTRs. Families with few or no tandem arrays (e.g. TRIM-65) as well as those with higher tandem formation instances (e.g. TRIM-01 and TRIM-33) both exhibit large numbers of partial matches, suggesting that recombination-driven diversification is also common in non-autonomous families. Together, these results indicate that recombination in TRIM elements gives rise not only to tandemly-arranged copies and solo-LTRs, but also to a substantial number of structurally altered, partially conserved TRIM derivatives.

### Not all non-autonomous LTR retrotransposons have a traceable origin

The origin of non-autonomous families can be determined in two ways: (A) through the presence of residual protein domains derived from autonomous elements, and (B) through detectable sequence similarity to extant autonomous partners.

(A) Firstly, among the 115 families in our dataset, 43 families retain remnants of coding domains from autonomous LTR retrotransposons (Ty1-copia: 10; Ty3-gypsy: 33), as identified with DANTE [24]. Based on their domain content (Additional Tab. 1.1 and 2), these families fall into two categories:

i. 28 families contain only GAG/PROT domains. Also described as TR-GAGs in *Coffea* [55] and later in Arabidopsis and the Triticaceae [56,57];
ii. 15 families retain at least one additional, partly decayed POL-associated domain, such as reverse transcriptase (RT), RNAseH (RH), Integrase (INT) and chromodomains (CHD). Eight of these families also carry GAG/PROT remnants, whereas seven lack them.

Notably, 24 out of 43 domain-carrying families show no detectable similarity to any recent autonomous LTR retrotransposon in our sugar beet specific database [34]. Within these families, the number of copies preserving fragmented domains varies considerably. In eight families, every identified copy retains coding remnants; all eight are small families with five or fewer full-length elements. More often, domain preservation is patchy: some family members retain coding fragments, whereas others display the typical non-autonomous structure with no remaining coding capacity.

(B) A second line of evidence to trace TRIM origin comes from the dependence of non-autonomous elements on the transposition machinery of autonomous retrotransposons. It is often assumed these elements establish a partnership with a corresponding autonomous element, which enables mobilization, although the molecular mechanisms underlying such partnerships remain unclear. Shared sequence similarity between autonomous and non-autonomous retrotransposons is often suggested to direct these partnerships, which also possibly enables allocation of non-autonomous families to either Ty1-*copia* or Ty3-*gypsy* superclade.

In our dataset, 30 non-autonomous families show a direct relationship to autonomous LTR retrotransposons. These relationships can be grouped into three levels of similarity:

i. high LTR similarity of non-autonomous/autonomous LTRs of 70% or higher (n=10);
ii. moderate LTR similarity (50-70%) between non-autonomous and autonomous elements (n=12); and
iii. low LTR similarity (<50%), but with supporting evidence from similarity in internal regions (often fragmented) or from the presence of 5′ UTRs and/or coding remnants such as GAG, PROT, or INT (8 families).

Taken together, the origin of non-autonomous elements can be reconstructed either through preserved coding domain remnants or through sequence similarity to known autonomous elements. In our dataset of 115 families, we were able to determine 24 families carrying reminiscent of coding capacity and 11 families with a similarity to an already described autonomous sugar beet LTR retrotransposons only, while 19 families exhibit both. In summary 54 of 115 families support the origins explained by (A) and (B). Especially for the last group we attempted to identify microduplications that might mark breakpoints associated with loss of coding capacity. However, no consistent signal was detected. Instead, non-autonomous elements frequently contain multiple short duplicated sequences within their internal regions (Additional Fig. 4), obscuring clear breakpoint signatures.

The general proportion of both autonomous LTR retrotransposons superfamilies across the sugar beet genome is almost even [34]. In our dataset, across all traceable non-autonomous families, the distribution between superfamilies is strongly skewed: Ty1-*copia* (12 families) versus Ty3-*gypsy* (42 families), corresponding to an approximate ratio of 1 : 3.5.

### Non-autonomous LTR-retrotransposons are scattered across the genome

Genomic localization of repeats within the genome may be associated with conservation and genomic behavior. Therefore, we analyzed the chromosomal positions of all full-length TRIMs in our dataset within the assembled pseudochromosomes of the *Beta vulgaris* sugar beet reference genome (Fig. 5). We further compared TRIM distribution to that of detected full length sequences of autonomous LTR retrotransposons in our reference database.

**Fig. 5:**
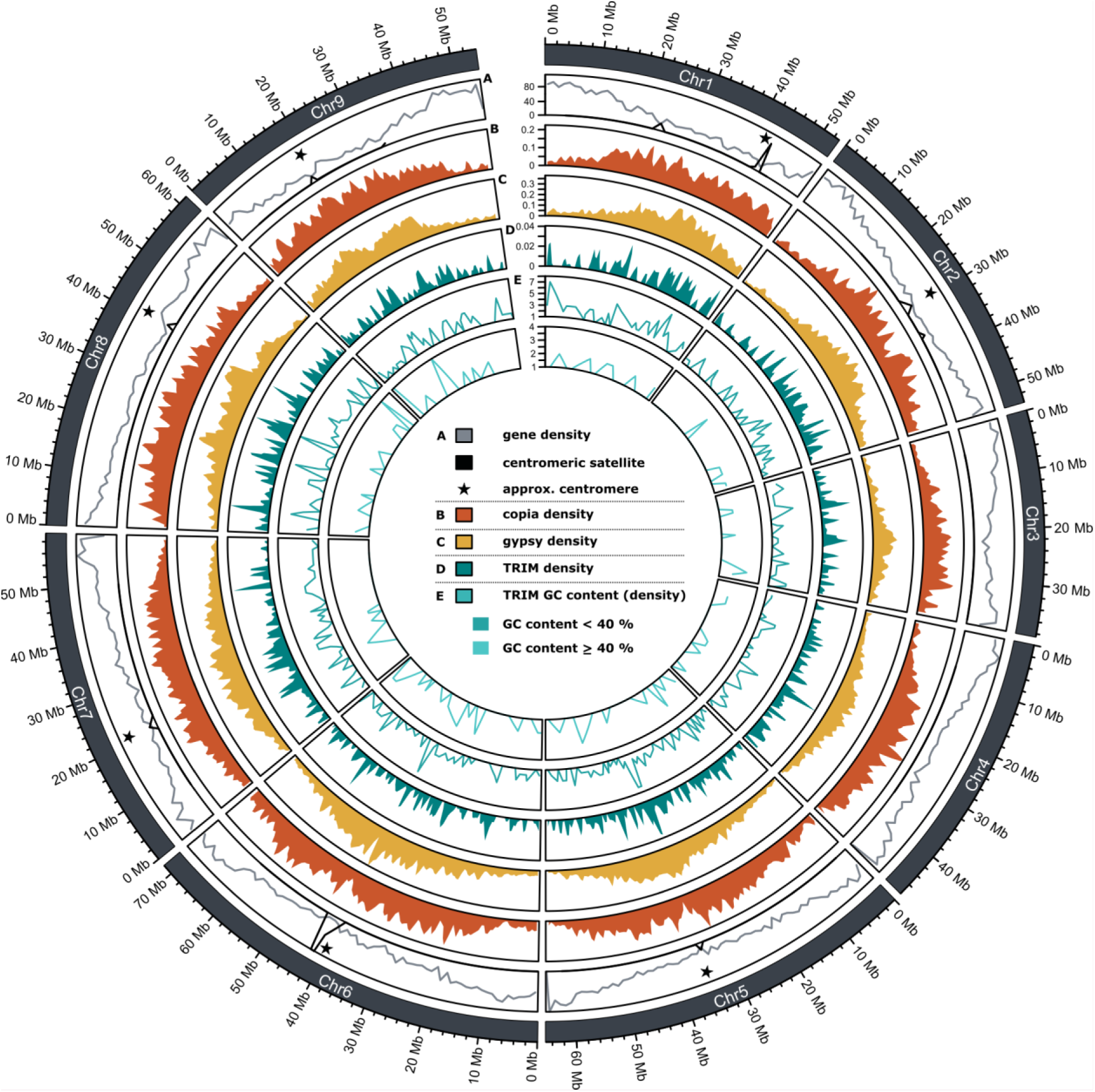
Localization of TRIM full length sequences across sugar beet pseudochromosomes. The density of autonomous LTR retrotransposons as well as non-autonomous elements (TRIMs) are shown as density plots with a window size of 1 Mb and are colored according to the superclade (orange = Ty1-*copia*, yellow = Ty3-*gypsy*, blue = TRIM). We furthermore added a track for the GC content of all full length sequences within our dataset, which are shown in a lighter teal and are into two tracks for sequences with less or more than 40%.

TRIM density is strongly reduced in centromeric regions (Fig. 5, asterisk; indicated by centromeric satellite repeats and Ty3-*gypsy* accumulation). Instead, non-autonomous elements reside in gene- and Ty1-*copia* rich regions, especially along chromosome arms. To complement this analysis, we examined the G/C-content of all non-autonomous sequences in our dataset, which ranged between 23% up to 53% G/C. Both groups tend to show distinct spatial distributions, with G/C-rich copies tending to appear in more gene-dense regions.

To comprehend the analysis, we performed a gene association study between a published gene annotation [39] and our dataset. We identified non-autonomous retrotransposons from our dataset associated with genes either by direct overlap with gene bodies (n=216) or by proximity within 5 kb flanking regions. Among gene overlaps, the vast majority were intronic (n=156; 72%), whereas only a small fraction overlapped with exonic or coding regions (n=60; 28%), potentially relevant for alternative splicing. We observed 627 instances of non-autonomous element insertions located close to genes within 5kb flanking regions. Gene flanking elements showed a distribution of 51% (n=315) upstream and 49% (n=312) downstream localisation in a strand-aware manner (additional Table 3B). Notably, individual copy insertions were frequently associated with multiple neighboring genes (Additional Table 3A), reflecting their location within potential shared regulatory hotspots. The enrichment of non-autonomous LTR retrotransposons in upstream promoter-proximal regions supports a potential regulatory role for these elements, either through the provision of cis-regulatory sequences or via broader effects on local chromatin structure.

## Discussion

Here, we provide a detailed overview about the challenging *de novo* identification of non-autonomous LTR retrotransposons and provide insights into the highly changing non-autonomous LTR retrotransposon landscape.

### Identification of non-autonomous LTR retrotransposons requires finetuning Non-autonomous

LTR retrotransposons are structurally minimalistic and highly variable, making them among the most difficult repetitive elements to identify in plant genomes. Their lack of conserved protein-coding domains, broad size spectrum, and lineage-specific sequence divergence mean that conventional *de novo* LTR retrotransposon pipelines systematically underdetect them or classify them as fragmented autonomous elements. Consequently, the true diversity and genomic impact of non-autonomous LTR retrotransposons remain substantially underestimated across plant genomes.

Detection of transposable elements in general and non-autonomous sequences in particular, relies on accurate boundary detection, and thus benefits from high quality genome assemblies and sequencing strategies [57–59]. Long-read sequencing is especially relevant for recently integrated elements; for example BARE2 elements with recent integration and highly similar LTR sequences being not represented or suffering from gaps in short read based genome assemblies [56]. Accordingly, we chose a Oxford Nanopore long-read genome assembly for a reference sugar beet genotype [39].

To capture the full spectrum of non-autonomous elements, from ultra-short TRIMs to unusually long LARDs, we employed a three-parameter LTR_Finder setup ranging from stringent to relaxed. Evaluation of our results (Additional Fig. 2) indicate that combining different parameter sets is not necessary. Hence, we recommend slight adjustments to the minimal LTR length and LTR distance parameters (-l and -d). Our data illustrates why pipelines using default LTR_Finder settings (including EDTA) miss substantial portions of the non-autonomous landscape.

After initial LTR_Finder analysis, we focused on generating a dataset of intact, full length sequences suitable for quantification. Unlike other pipelines, we targeted non-autonomous elements exclusively, performing a BLAST against the RepeatExplorer2 database to exclude sequences with coding potential from autonomous elements. This approach prevents misclassification: non-autonomous copies with partially deleted or missing coding regions are not automatically excluded. Furthermore, we adopted a semi-automated workflow combining high-throughput processing with dotplots, automatic feature detection and manual curation. This hybrid computational/manual approach was crucial to maintain accuracy: dotplot inspection remains one of the most robust ways to validate element structure (e.g. full length, tandemly-arranged copies, indel availability) and automated pattern recognition helped to scale this process to thousands of candidates.

We recommend an additional self-blasting after quantification, adapted from the DANTE-LTR approach, which removes highly truncated or single LTRs before dotplot analysis. An applied length threshold, excluding all intact copies with indels greater than 15% of the median LTR length within the family, is beneficial because otherwise larger insertion/deletions influence identity value calculation which could be misleading [56].

The resulting dataset of 1,581 intact (= correct LTR boarders, availability of PBS and PPT, no larger indels) non-autonomous LTR retrotransposons to our knowledge are one of the most comprehensively curated catalogs within one plant species. By using LTR sequences for clustering and comparing multiple strategies, we were able to define 115 TRIM families. Our clustering benchmarking results highlight why relying solely on high-throughput clustering tools is not beneficial for non-autonomous elements:

- All-against-all BLAST is recommended to detect short, high-identity matches. Additional Markov-clustering (MCL) performs best for capturing homologous groups without oversplitting.
- High-speed tools using k-mer based methods, such as MMseqs2 and CD-Hit, are not suitable for shorter, variable TRIMs because of their low k-mer complexity.
- Although SiLix performs reasonably, it tends to split families because clustering based on strict homolog intervals is not beneficial for a dataset of highly variable length sequences.

In the end, we applied a more relaxed threshold compared to the 80-80 coverage rule recommended by Wicker et al., 2007 [13]. Relaxed thresholds are justified for non-autonomous elements, which lack highly conserved protein domains, e.g. like RT, and are consistent with prior TE studies [60–62].

Clustering short repetitive sequences remains a major challenge, for multiple reasons: (1) Low sequence complexity of non-autonomous elements; and (2) high structural and length variability, complicating clear boundary delineation and subfamily identification. Manual curation and multiple sequence alignments remain essential for reconstructing family structures also within non-autonomous LTR retrotransposon landscapes [19,63–65]. Together, our findings underscore a central conclusion: accurate identification of non-autonomous LTR retrotransposons requires dedicated workflows.

### Non-autonomous LTR retrotransposon diversity is vastly underestimated

Our curated non-autonomous dataset reveals a remarkable continuum of structural variability. Family median lengths range from 290 bp up to 12 kb (42-fold difference) and LTR sizes vary from 72 bp to 5.5 kb (76-fold difference). Traditionally, non-autonomous TRIMs are defined as shorter than 1 kb [14]. Eight families in our dataset meet this strict definition and resemble typical TRIM representatives such as *Wukong* [66], Cassandra [18,32,67–69] or TRIMs from *Brassica*, apple and rice [70,71]. Extending the definition to 1.5 kb as introduced in Gao et al (2016) [17] captures 15 families total. Non-autonomous elements exceeding 4 kb in length, so-called LARDs [15,72] correspond to medium, long and extra long element sizes in our dataset. However, intermediate size ranges from 1.5 - 4 kb remain undefined, despite reports of “long-TRIMs” in rice with lengths of 1.6 and 5 kb [73].

Despite their structural variability, most families (96%) exhibit high pairwise LTR similarity above 85%, suggesting either recent mobilization or slow accumulation of mutations. Furthermore, we observe high TSD retention across our dataset, indicating young populations. This is in hand with rapid and recent integration of non-autonomous LTR retrotransposons within genomes in high or low copy numbers [74,75]. High LTR identities persist even in families with large LTRs, consistent with observations of longer LTRs showing higher LTR similarities [76]. Copy number, LTR identities and TSD retention rates, do not reliably allow conclusions regarding ages or activity: both high- and low-copy families exhibit high LTR identities, which indicates family-specific mobilization dynamics. Hence, our dataset is biased due to the length threshold we applied removing eroded copies, which favors conserved elements.

PBS and TIR motif analysis add further evidence of origin diversity and modularity. Methionine-tRNA-derived PBS motifs dominate the whole non-autonomous population, which is often associated with Ty1-*copia* and chromoviral Ty3-*gypsy* families, but we also identify less common motifs, likely derived from tyrosine-, histidine- and valine-tRNAs, which rarely occur in known autonomous plant LTR retrotransposons [12]. These atypical PBS types highlight that non-autonomous elements frequently emerge from multiple autonomous progenitor lineages, including lineages that may no longer be active or detectable in the reference genome. Additionally, we observe Ty3-*gypsy* associated aspartic acid-tRNA-derived PBS motifs for non-autonomous families of medium and large length groups, indicating Ty3-*gypsy* origin.

Similarly, TIR motifs do not cleanly map onto superfamily predictions. Horváth *et al*. (2024) [77] recently characterised TIRs across autonomous LTR retrotransposons and reported superfamily-linked tendencies: TGTA enriched in Ty1-*copia*, TGAT in Ty3-*gypsy*, and TGTT restricted to a small subset of families from both superfamilies. Although these are reflected to some extent, non-autonomous elements deviate from these patterns, displaying lineage mixing and motif distribution across the whole population with no clear connection to length. We explain this by fewer structural constraints on non-autonomous element boundaries, while retaining those minimal features necessary for host integration after divergence from their autonomous progenitors.

Collectively, these results indicate that non-autonomous LTR retrotransposons are not evolutionary relics of their autonomous siblings, but a spectrum of structurally flexible mobile genetic components with a decent amount of activity [14,78]. They retain the essential machinery for recognition and integration, while showing high amounts of structural flexibility. This sort of modularity could reflect an evolutionary strategy distinct from autonomous elements: structural reduction enables tolerance, mobility and persistence enabling occupation of genomic niches inaccessible to larger retrotransposons.

### Recombination is a vital part of non-autonomous LTR retrotransposon biology

Recombination is a central mechanism shaping LTR retrotransposon landscapes. Our data demonstrates that non-autonomous elements are strongly influenced by recombination-driven processes. They display a dynamic turnover, which reflects their reduced structural constraints and their reliance on a few conserved motifs needed for host-dependent propagation.

Two major recombination routes have been laid out before: (1) illegitimate recombination, which generates truncated elements [79,80] and (2) unequal homologous recombination between LTRs of either the same or distinct elements, producing solo-LTRs or tandemly-arranged copies [79]. These recombination events are very common in plants [81,82] and our non-autonomous dataset provides a quantitative view of both pathways.

Solo-LTRs are detectable in 102 of 115 families, with solo/full-length ratios spanning four orders of magnitude, from nearly absent to more than 100 : 1. Presence of solo-LTRs in families with high LTR identity values suggest recombination happens continuously, even upon young populations and is not restricted to old, accumulated copies. As TSD retention rate for solo-LTRs is only 49%, solo-LTR formation is favored by inter-element unequal recombination. Also tandemly-arranged copies are widespread across the non-autonomous landscape in the sugar beet genome, occurring in 38 families and spanning all length and LTR sizes. Arrangement in tandems is well-described for Cassandra elements and other TRIMs [14,16,18,83]. 71% of elements in tandem retain TSDs, indicating that they originate from intra-element recombination. Although being less represented, formation of TA copies from ectopic recombination is present in sugar beet and likely contributes to sugar beet genome plasticity [84]. Tandem formation occurs independently of copy number, LTR identity or element size, contradicting earlier observations from soybean where tandemly-arranged elements are enriched among LTR retrotransposons with larger LTRs [85]. In contrast, solo-LTR formation shows no clear relationship to size, copy number or identity values, aligning with LTR retrotransposon studies in conifers [53]. Our results indicate that recombination is acting on a wide length spectrum and even very small elements, like TRIM-01, can participate in tandem formation in sugar beet.

So, as length, copy number and LTR identities are not driving recombination, what is? The size ratio of full length/LTR has been proposed to facilitate tandem formation [86], and also in our dataset, we observe a tendency regarding this factor: nine families with ratios larger than five lack detectable TA copies.

Besides locus-sensitive recombination, we also observe evidence for illegitimate recombination, such as truncated or hits suffering from indels, and modular re-shuffling (additional tables 1). Template switching during reverse transcription is a well-established driver of TE innovation [11,87–93]. Template switching has also been documented in autonomous LTR-RTs in sugar beet [36] and similar modular recombination is evident in several non-autonomous families in our dataset. Such hybrid elements illustrate how recombination can reshuffle structural components across families, generating novel combinations of LTRs and internal regions [9,94]. Such events are not limited to families with shared sequence ancestry: chimeric elements involving highly divergent families emphasize that recombination-mediated innovation is a genome-wide process rather than a lineage-specific mechanism.

Taken together, we determine that non-autonomous elements do not represent a “dead end” of LTR retrotransposon evolution. Instead, recombination, both homologous and illegitimate, is a major driver of structural diversification in non-autonomous elements. This contributes directly to the dynamics of repeat landscapes in sugar beet and shows that recombination is not merely a decay mechanism but facilitating structural reinvention and diversification.

### Non-autonomous and autonomous TEs form non-similarity based complex partnerships

Non-autonomous elements without (complete) coding capacity require the enzymatic machinery of autonomous elements. Such cross-mobilisation is well established for other non-autonomous transposable elements, such as (Alu) SINEs [95,96], MITEs [97], and even endogenous retroviruses [98] and also suggested for non-autonomous LTR retrotransposons [14]. For instance cross-mobilisation has been shown for a certain type of non-autonomous LTR retrotransposons, so called retrozymes, which rely on Ty3-*gypsy* reverse-transcription machinery through partial sequence similarity [99]. Yet the nature of these partnerships and their specificity remain poorly understood. Our dataset provides new insights into the mobilization mechanisms of non-autonomous LTR retrotransposons, revealing a spectrum of possible interactions ranging from strict similarity pairing [17] to opportunistic or generalistic mobilization.

In our dataset, 30 families exhibit obvious similarity to known autonomous LTR retrotransposons; either via shared LTR or internal sequences, or both. These relationships span both Ty1-*copia* and Ty3-*gypsy* again supporting repeatedly and independently emergence of non-autonomous elements across multiple autonomous lineages. Nevertheless, high LTR identities with autonomous elements are only observable in 10 families, emphasizing that many autonomous progenitors may no longer be present in the current reference genome and mobilization is happening in a more generalistic manner.

Protein domain remnants further highlight the complexity of non-autonomous divergence and origin. We detect coding motifs in 43 families within our dataset, identified with DANTE. Yet only 51% of these families show consistent motif availability across all members. In the remaining families, domain presence appears patchy and uneven and is similar to structural variation of autonomous LTR retrotransposons in plants [56]. Patchy variability indicates domain loss and partial decay can occur gradually and independently across families, probably driven by small internal deletions, reverse transcription errors and recombination. Similar patterns underlie formation of TR-GAG elements in *Coffea* [55] and Arabidopsis [57]. A well-studied case of such interaction is the RLG_Quinta/RLG_Cereba pair in wheat, where non-autonomous RLG-Quinta contributes structural GAG but relies on autonomous RLG_Cereba for activation, forming a mutualistic partnership [100]. In contrast, BARE-2 in barley is a defective chimera lacking a functional GAG domain and depends on its autonomous partner BARE-1 for packaging, representing inter-element parasitism.

Integration and mobilization of LTR retrotransposons without *integrase* was recently shown in yeast [101] and indicates active transposition of “defective” elements is a common process across kingdoms. Most likely, plant LTR retrotransposons similarly engage in cross-activation: autonomous elements can mobilize non-autonomous copies even with minimal shared sequence identity [56]. Our dataset supports this assumption with 61 families lacking any detectable similarity to autonomous elements yet retaining high LTR identities and TSDs. These families must rely on generalist mobilization, where active autonomous elements provide the machinery for *trans* interaction [6,14,102].

PBS and TIR motifs, often considered pointing to superfamily origin, reinforce this assumption. While many non-autonomous elements carry PBS-Met motifs characteristic to Ty1-*copia* and chromoviral Ty3-*gypsy* elements, rare PBS types and mixed TIR features suggest repeated emergence from multiple autonomous elements, followed by subsequent divergence. The inconsistent presence of PBS/TIR motifs and LTR structural features further indicates that mobilization is flexible, not constrained to strict element-specific compatibility and rather working as a network.

Altogether, our observations support the assumption that non-autonomous LTR retrotransposons are mobilized in a decentralized way, using the enzymatic machinery, which is available at this time and space. This also explains how diverse and re-shuffled non-autonomous elements can proliferate and persist despite lacking the machinery for autonomous propagation.

### Same, but different: non-autonomous LTR retrotransposons mirror the autonomous elements’ behavior to their benefit

Distribution of non-autonomous LTR retrotransposons on sugar beet pseudochromosomes reveals a pattern: these elements are depleted in pericentromeric and centromeric regions, where heterochromatin and Ty3-*gypsy* elements accumulate. Instead, they are enriched along chromosome arms, particularly in gene-dense and Ty1-*copia* rich regions. This mirrors observations in several plant taxa including *Brassica species* [71], apple [70] and rice [17], in whose genomes TRIMs preferentially occupy euchromatic or moderately accessible chromatin.

This distribution is most likely biased by several biological factors: 1.) Size-based tolerance of non-autonomous elements in gene-rich regions by the host [14]. Most non-autonomous elements, because of their compact architectures, impose lower mutational and regulatory burdens compared to larger autonomous retrotransposons. This neutral behavior makes their insertion more tolerable in gene-dense regions, which are otherwise under strong selection against repeated insertions. For example, smaller genes of less than 2 kb with conserved association of small TEs remain more expression-stable than those next to larger TEs [103]. 2.) Epigenetic regulation promotes open, accessible chromatin [104]. Encounter rates between non-autonomous LTR retrotransposons and autonomous protein machinery is more likely in euchromatic regions, which enable transcriptional access through the host. Enrichment of non-autonomous elements in these regions may therefore reflect a permissive integration hotspot and simultaneously enhanced conditions for mobilization. 3.) Non-autonomous elements prefer Ty1-*copia* enriched regions. In our dataset, non-autonomous elements mirror the genomic landscape of Ty1-*copia* rather than Ty3-*gypsy* elements, even though feature analysis suggests many families originated from Ty3-*gypsy* progenitors. This indicates that genomic distribution is decoupled from evolutionary origin; non-autonomous retrotransposons prefer integration with accessible chromatin and not necessarily where their progenitors resided.

GC-content is also an indicator for genomic niche occupation. Usually, GC contents of plants vary widely and are species-specific [104]. Non-autonomous elements in our dataset span a wide range of GC-contents between 23 - 53%. We observe localization of GC-richer families in gene-dense regions, which is consistent with TE-gene association patterns observed for autonomous elements such as *Tos17* in rice [106]). However in contrast to GC-rich and massively expanding Huck elements in maize [107], GC content alone is not an indicator for proliferation in sugar beet: high copy number families occur across both GC-poor and GC-rich families and families with or without a designated autonomous partner. This reinforces that non-autonomous element distribution across the genome is happening by preferred integration into GC-richer regions rather than GC-driven expansion.

We identified 216 TRIM-associated genes, with the majority of insertions intronic (72%). Exonic TRIMs, although less frequent, illustrate that small elements can integrate into coding regions without necessarily terminating gene function. This could also be explained by smaller TEs near genes that exhibit less disruptive effects and greater expression stability than large TEs [103].

These patterns collectively support that non-autonomous LTR retrotransposons populate within the genome in an opportunistic way. They prefer open chromatin for integration and mobilization, preferably in gene-rich regions. Because of their size-related smaller genomic imprint, these niches may be more suitable for non-autonomous elements.

### Non-autonomous elements are modules of genomic plasticity

Beyond shaping genomic architecture through copy number variation and recombination, non-autonomous elements contribute to the structural composition of plant genomes. Non-autonomous elements are structured in a modular way: 1.) structural modules - LTRs; necessary for transcription start, 2.) replication module - PBS and PPT, needed for reverse transcription of the RNA and occasionally 3.) remnant protein domain modules. This makes them potent contributors to genomic innovation.

In our dataset, we observe that these modules evolve in a semi-independent way: reductions of coding domains decrease coding potential, but LTRs still enable structural integrity and activation. Recombination of these structural features can exchange LTRs without disrupting mobility and template switching can generate chimeric elements with mixed ancestry across different modules. This architecture mirrors mechanisms described for autonomous elements, where recombination and domain mixing create new retrotransposon lineages [87–89]. Non-autonomous elements thus participate fully in the modular evolution of plant genomes despite their reduced structure.

Furthermore, LTRs are hotspots for regulatory motifs, also in TRIMs. Several TRIM lineages such as Cassandra and Helenus/Ajax even obtained rDNA or tRNA-like motifs which enabled stress-independent transcription [18,32,67,108]. Additionally, LTRs of non-autonomous elements contain promoter/terminator-like structures which could act as alternative promoters or terminators of nearby genes and TE-derived transcription factor binding sites are most likely to influence gene expression [109]. Because non-autonomous elements prefer gene-rich regions their regulatory potential most likely affects gene expression. Especially for smaller size families this allows persistence without strong epigenetic silencing.

Structural compactness and formation of tandemly-arranged elements as well as observations of non-autonomous elements harboring repetitive regions suggest a role in formation of satellite DNA-like repeats. Similar processes have been documented for other plant TEs [110]. Thus, in addition to providing potential regulatory elements, non-autonomous elements may also be drivers of structural innovation.

Taken together, the genomic distribution and association with genes of non-autonomous elements reflect selective forces of non-autonomous TE modules: 1.) Non-autonomous TEs may persist in genic regions; 2.) insertions may not trigger epigenetic silencing of nearby genes; 3.) LTRs can co-opt as new regulatory elements; 4.) Structural turnover generates recombination-driven ongoing novelty.

In this framework, non-autonomous LTR retrotransposons are not leftovers of autonomous decay but active participants in gene-rich chromosomal landscapes.

### Solving the dilemma of classification: How to name a TRIM, LARD, TR-GAG?

Despite two decades of research, terminology for non-autonomous LTR retrotransposons remains fractured. “TRIMs”, “long-TRIMs”, “SMARTs”, “LARDs”, “TR-GAGs”, and “defective” LTR retrotransposons all describe subsets of non-autonomous retroelements, but the boundaries between these categories blur when examined at scale. Our sugar beet dataset illustrates why classification is problematic.

Size thresholds do not map biological reality. Originally described as “TRIMs” or “SMRTs”, small non-autonomous elements were defined as less than 1 kb in size [14,111]. In our dataset this applies to 8 out of 115 families. If we extend the definition to less than 1.5 kb in size as proposed in Gao et al., 2016 [17] the threshold would still eliminate more than 85% of families in our dataset. Elements larger than 4kb in size or so called LARDs [15] encompass all elements of medium sizes and above in our dataset. But no terminology exists for size ranges from 1.5 - 4 kb size ranges, yet this group contains many trustworthy non-autonomous families in the sugar beet genome. Thus, this terminology represents the extremes of a continuum, not discrete types.

Coding-capacity categories are equally ambiguous. The discovery of TR-GAG elements [55] introduced a category of non-autonomous retroelements that retain a GAG ORF while lacking enzymes with a length of less than 4 kb assumed to be “semi-autonomous” [56]. However, our dataset includes 29 families with GAG/PROT domains, but these exceed the proposed size spectrum. A further 14 families contain degraded fragments of RT, RH, or INT but lack complete coding capacity necessary for self-driven mobilization. Although occasionally described by authors within repeat landscapes of various plant species no terminology exists for these kinds of elements (yet). Thus, domain remnants serve as clues of evolutionary history but not as robust classification anchors.

Only a fraction of non-autonomous elements has a traceable origin. Only 30 of 115 families share detectable similarity to known autonomous elements in sugar beet, and only 10 of these can be assigned with >70% LTR identity. Absence of similarity is most likely influenced by recombination-mediated remodeling or lineage extinction of the former autonomous progenitor. Consequently, origin cannot be reliably interfered with for most non-autonomous elements, even though their high LTR identities and TSD retention rates demonstrate recent activity.

None of the detected features in our dataset (PBS, PPT, TIR) alone allows classification. Furthermore, great size variety and variable TIR motifs do not enable architecture dependent superfamily association.

Together these patterns show that non-autonomous LTR retrotransposons cannot be reliably assigned into the recent terminology. We propose a conceptual shift: instead of “non-autonomous” equating “short and non-coding”, we advocate using the term “non-autonomous” for elements with non-functional transposition, regardless of size and presence of fragmented ORFs. We therefore propose keeping the name TRIM as a classification system for non-autonomous elements and introduce them as a third superfamily next to Ty1-*copia* and Ty3-*gypsy*. This follows the classification logic already used for other non-autonomous TE orders, such as SINEs and MITEs. We furthermore advocate for keeping definitions such as TR-GAG, Cassandra, or retrozymes to describe elements with designated features in a lineage-like way.

## Conclusion

Non-autonomous LTR retrotransposons represent a flexible and recombination-driven type of transposable elements, which evolve under relaxed constraints while preserving minimal motifs required for integration and mobilization. *De novo* identification of these elements depends on good quality genome assemblies as well as fine-tuned parameter settings and manual curation. Our data provides evidence for repeated emergence of non-autonomous elements from diverse autonomous progenitors. They undergo rapid structural remodeling through recombination and integrate preferentially into gene-rich chromatin. We propose non-autonomous elements persist due to their size and low regulatory burden in these regions and contribute to genome innovation by providing regulatory and structural features. Mobilization of non-autonomous elements is not dependent on fixed partnership and rather happening in a generalized way. Thus, non-autonomous elements are not static remnants but active, evolutionary, dynamic participants in shaping plant genome architecture.

## Declarations

### Ethics approval and consent to participate

Not applicable.

### Consent for publication

Not applicable

### Availability of data and materials

All identified non-autonomous full length sequences as well as corresponding solo-LTRs and tandemly-arranged copies are publicly available on Zenodo: https://doi.org/10.5281/zenodo.18174724.

Reference sequences for each TRIM family are available in the BeetRepeats database ([34] https://zenodo.org/records/8255813).

Furthermore, detailed workflow description as well as code for file processing is stored under github: https://github.com/TRIMpossible/TRIM_identification.

We used large language models ChatGPT and GPT5 to facilitate the scripts provided on github in order to improve user accessibility.

The classifier workflow to recognize full length copy patterns from flexidot output dotplots is available in github as well: https://github.com/tudipffmgt/Flexidot-Classifier.

### Authors’ contributions

Analysis: SM

Software, Code Scripting: SM, FM

Writing - draft: SM

Writing - review and editing: all

Figures: SM, FM

Conception: SM, TH

## Acknowledgements

We thank Ludwig Mann for excellent scientific and bioinformatic advice during the preparation of this study.

We thank Nicola Schmidt for curation of the BeetRepeat database and inclusion of TRIM reference sequences.

