## Supplement Figures for "Hide and seek: *de novo* identification in sugar beet reveals impact of non-autonomous LTR retrotransposons"

**Additional figure 1: Detailed overview of the flexidot computer vision approach.**
Stepwise illustration of the automatically repeat structure classification from dotplot images.

*1. Detection of plots*
Detection of candidates requires the inspection of every single plot on every page, which are extracted in the binary image generated using Otsu thresholding (Otsu, 1979) followed by a contour detection algorithm. As the plot is of fixed size, only the contours larger than 100 x 100 pixels and with exactly four sides are kept resulting in 20 plots per page. Finally, those are sorted by their top-left corner, so that the plot order of flexidot is kept and the mask of every plot is stored for later visualization. For easier detection of possible LTR retroelements by means of image analysis, “noise” in every plot is removed using the morphological operation opening and a subsequent dilation with a 5x5 kernel. Again, for faster processing every resulting plot is cropped from the PNG file and stored separately.

*2. Detection of repeated regions*
Supporting the visual inspection of all plots resulting from flexidot, the challenge is to identify LTR retrotransposons. Furthermore, we added an optional setting allowing the identification of tandemly arranged elements and arrays. Therefore, every plot is iteratively imported and analyzed using several conditions. Initially, the two largest contours are extracted. If the contours are of significant size and appear both on top right and bottom left of the plot, then it is of high probability that a TE is present, else the plot is discarded from further analysis. Continuing to identify the arrangement, it is checked whether the largest five contours in the plot run from top-left to bottom right. If this is true, then both tandem elements and satellite elements are possible. If even more contours (< 6) in the plot are positioned diagonally from top-left to bottom-right, the tool identifies the plots as possible satellite elements, else as tandem elements.

*3. Visualization and annotation of internal TE domains*
As the mask and the plot number are stored according to the flexidot annotation, it is then possible to draw a coloured frame around every plot that might be a TE and all respective sequence numbers are stored in separate text files for further extraction of sequences from the input fasta file.
This workflow allowed quick and automated file processing instead of time consuming manual curation or further file processing

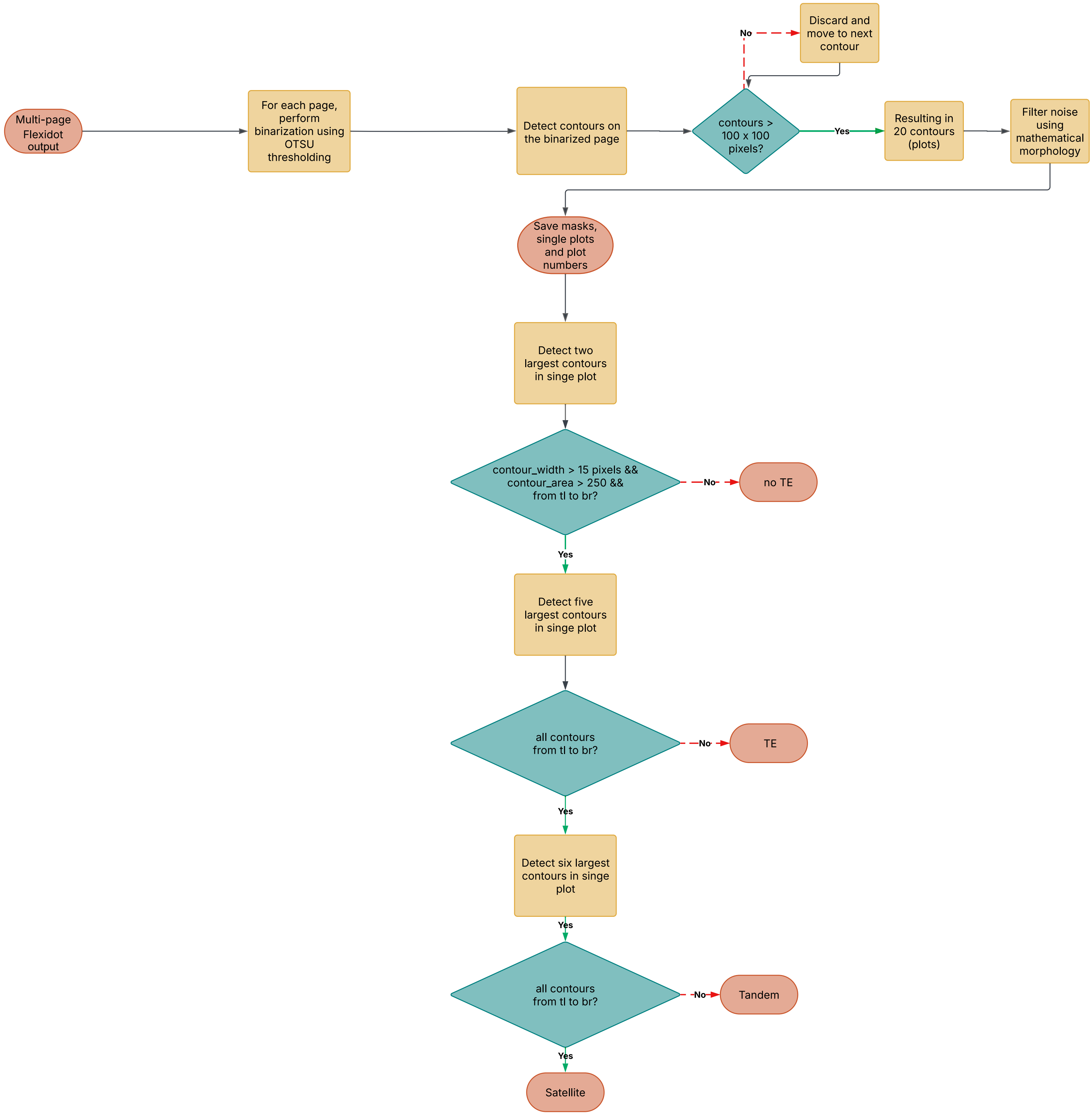

**Additional figure 2: Evaluation of de novo identification with LTR_Finder.**

LTR_Finder analysis were performed using three different parameter sets corresponding to the length of targeted elements. For each set, we retrieved length distributions corresponding to the chosen length parameters (A).

Visualized LTR_Finder score distributions confirm the majority of hits having low scores, as expected for non autonomous elements lacking coding regions (B).

We also provide the initial number of hits retrieved with LTR_Finder and their reduction during filtering, aswell as the final number of reference sequences used for quantification (C).

After quantification and condensation of full length sequences into families we traced how many families were identified per run (D).

The median length distribution of identified TRIM families reflects the correlation between overall lengths and initial chosen length parameters for the LTR_Finder analysis. Additionaly, 17 families were not detected by LTR-Finder but during quantification because of similarities shared with used reference sequences (E).

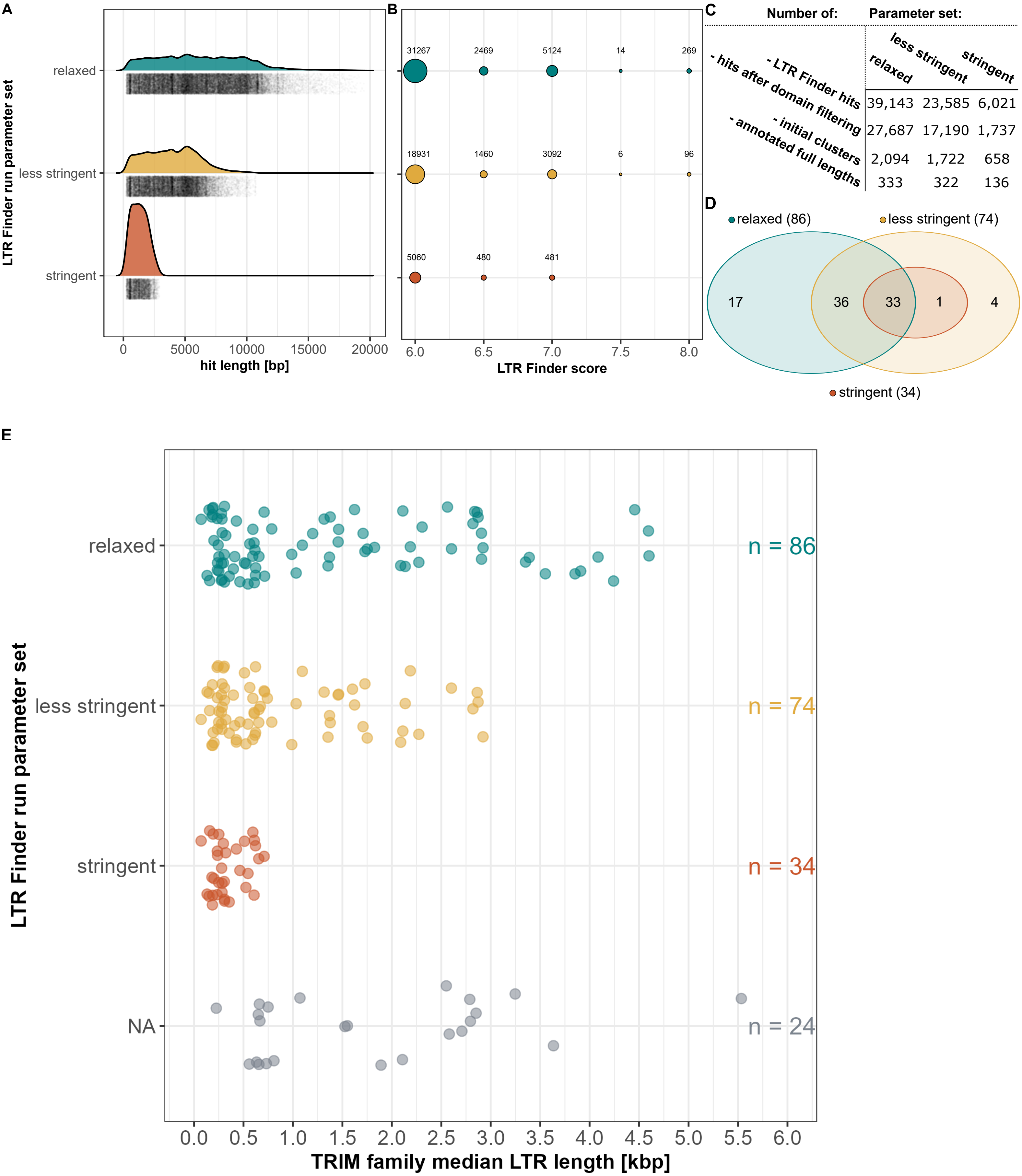

**Additional figure 3: Clustering tool and parameter test evaluation.**

As clustering tools are often used for coding or conserved protein and/or nucleotide sequences we performed parameter tests checking which is suitable for non coding and highly variable sequences such as non autonomous LTR retrotransposons. Two different clustering principles were used: kmer and similarity-network based tools. Shown in colours are always the number of resulting clusters based on the cluster forming controlling treshold values. Single sequence clusters for each parameter test are shown in grey. As input for all clustering approaches we used the dataset of 7,182 annotated full length sequences.

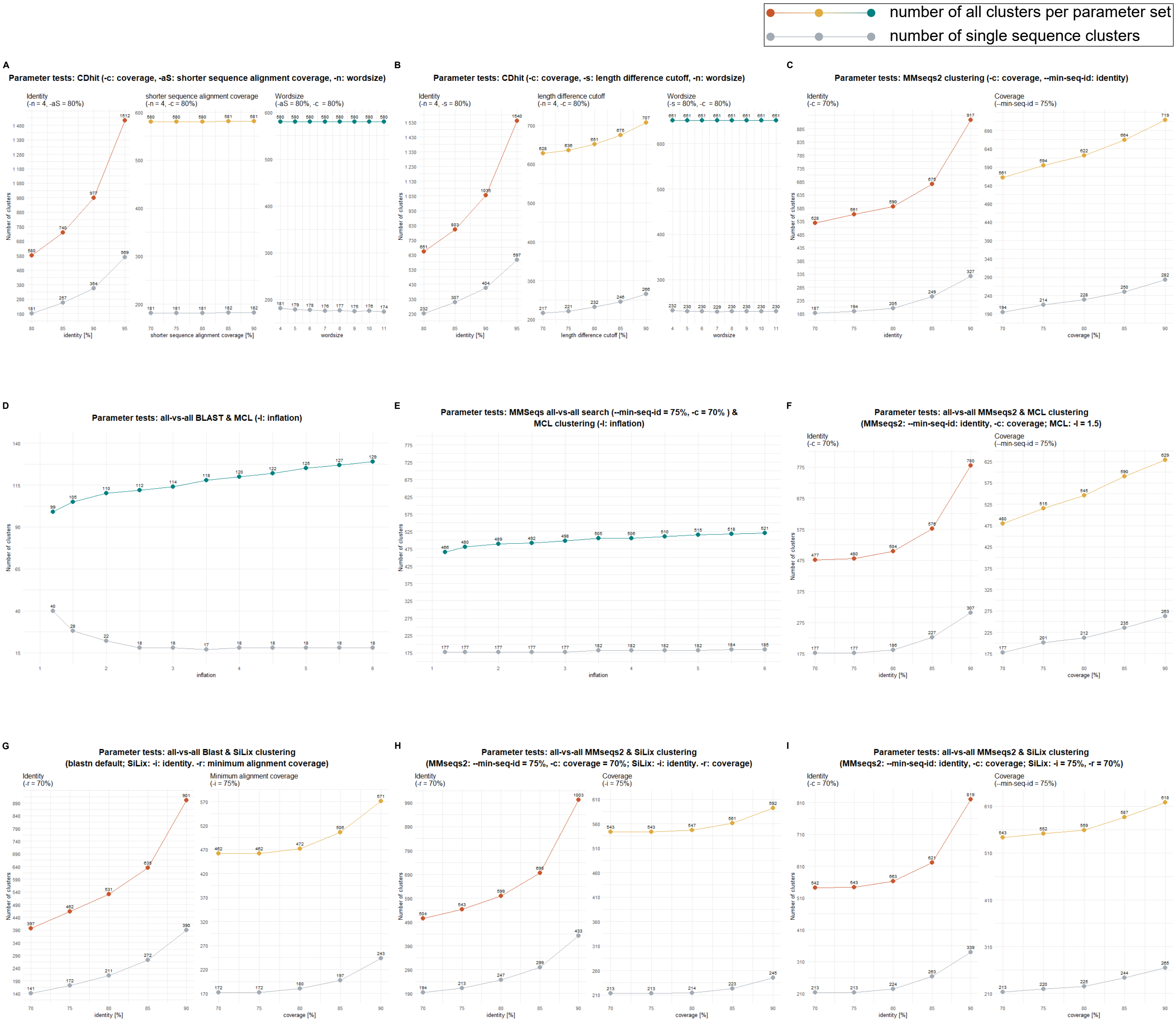

**Additional figure 4: Dotplot vizualisation of TRIM internal sequences.**

Shown are the internal regions of all TRIM reference sequences. Dotplots were created with flexidot with a wordsize of 7 and enabled reverse complement matching. All TRIM families show internal microduplication, which could be a result of illegitimate recombination (especially by microhomology-mediated end joining). Additionally, some families show accumulations of internal smaller stelllite like arrays.

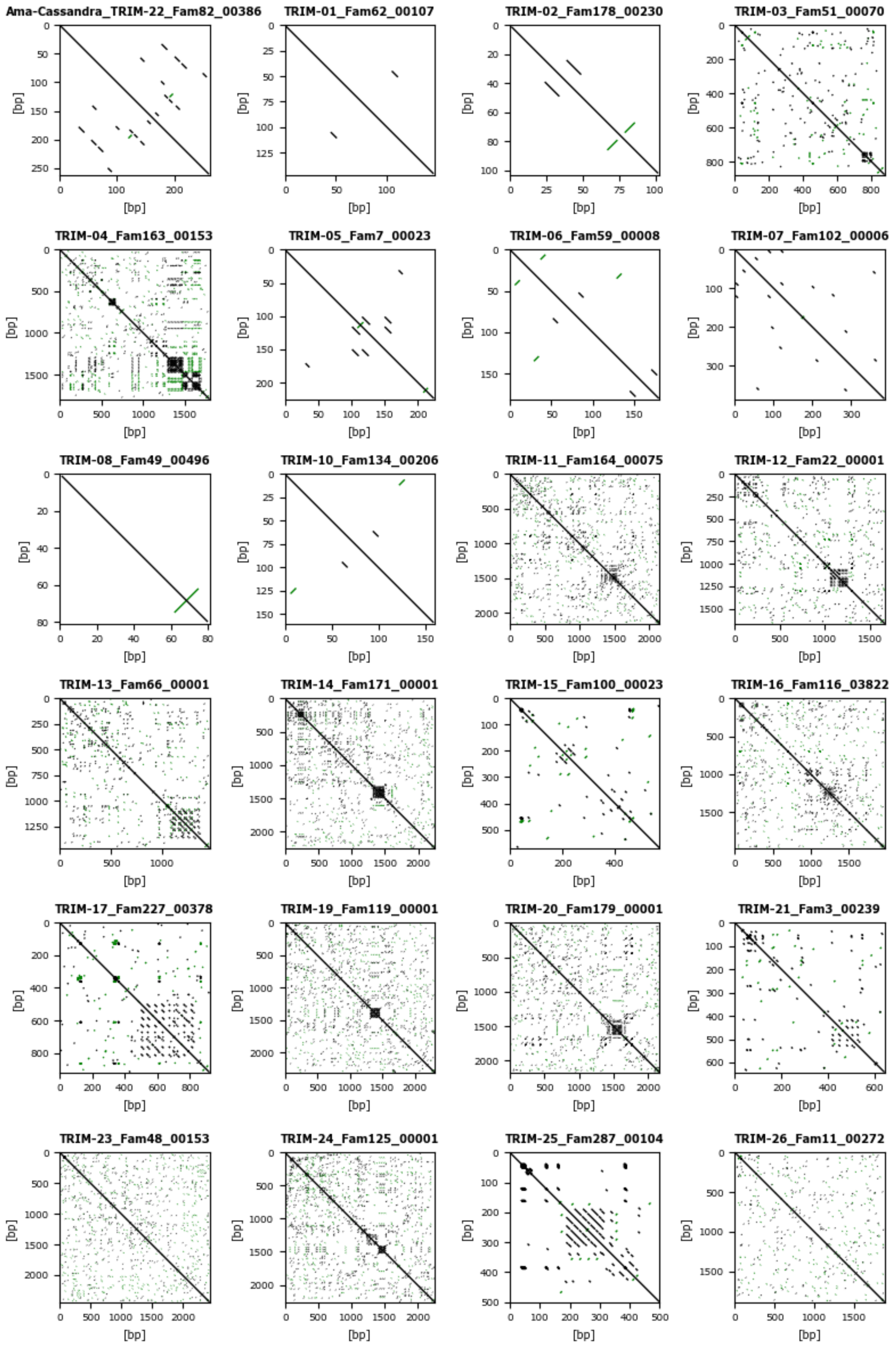

**Additional figure 4: Dotplot vizualisation of TRIM internal sequences.**
Continuation.

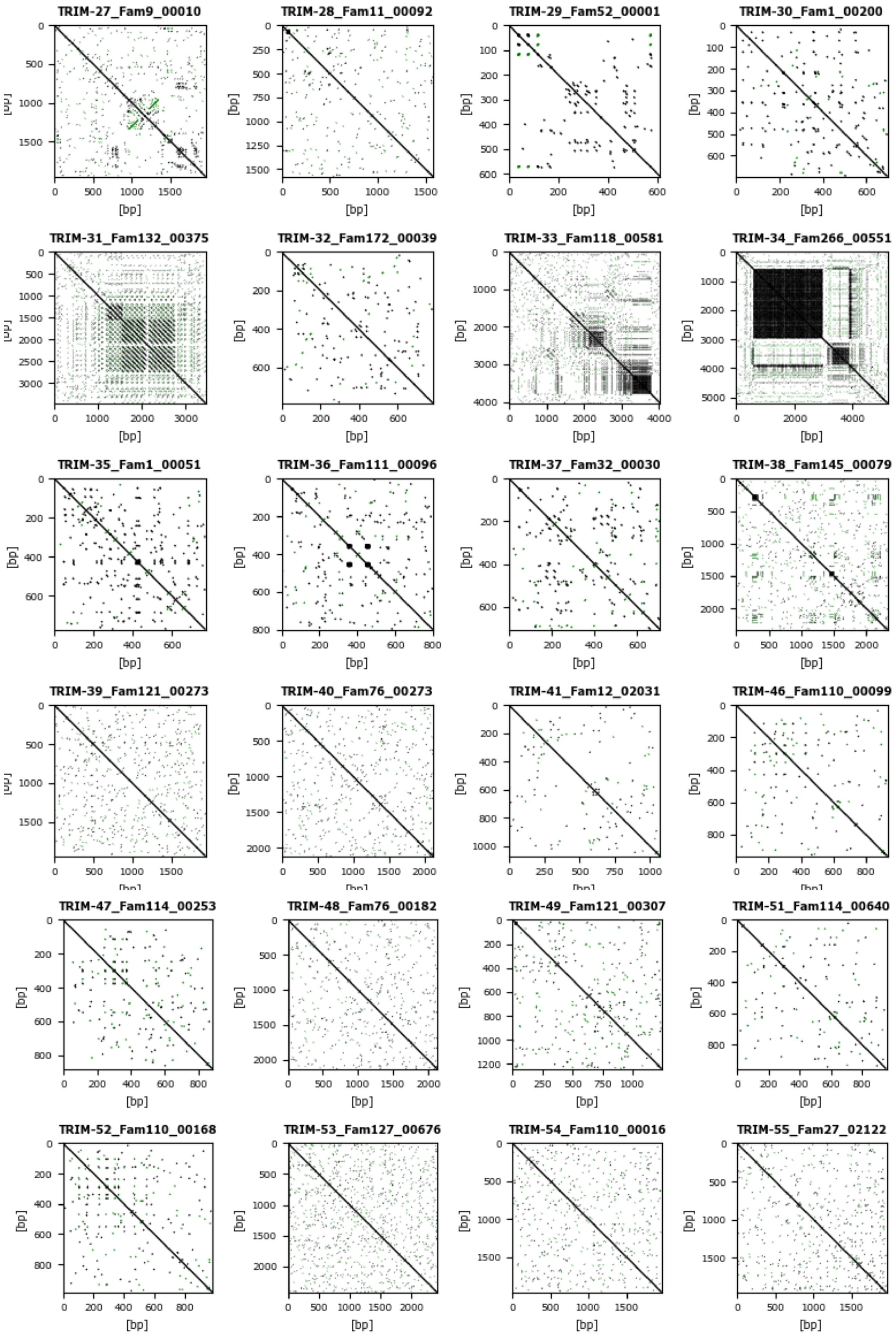

**Additional figure 4: Dotplot vizualisation of TRIM internal sequences.**
Continuation.

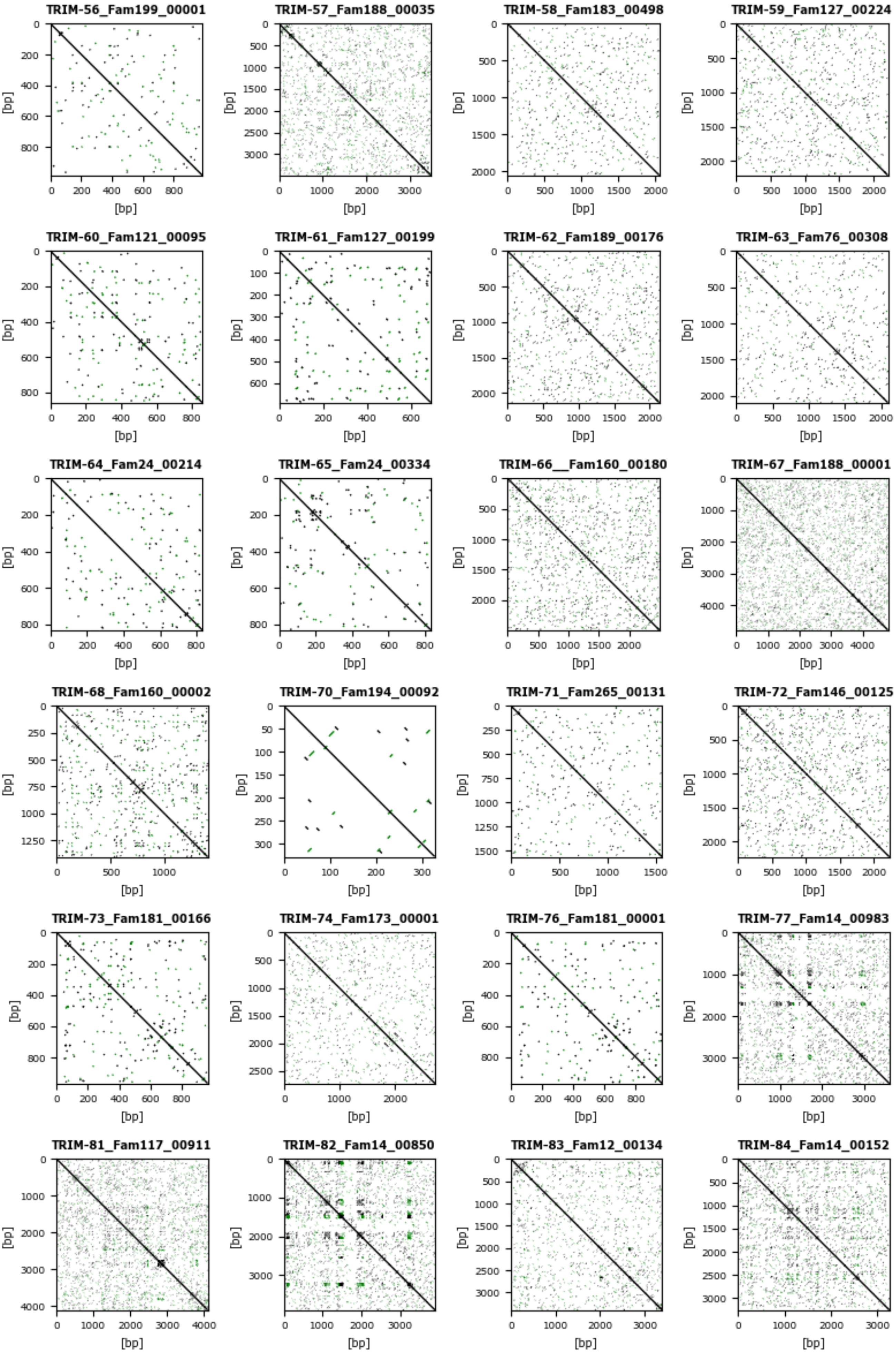

**Additional figure 4: Dotplot vizualisation of TRIM internal sequences.**
Continuation.

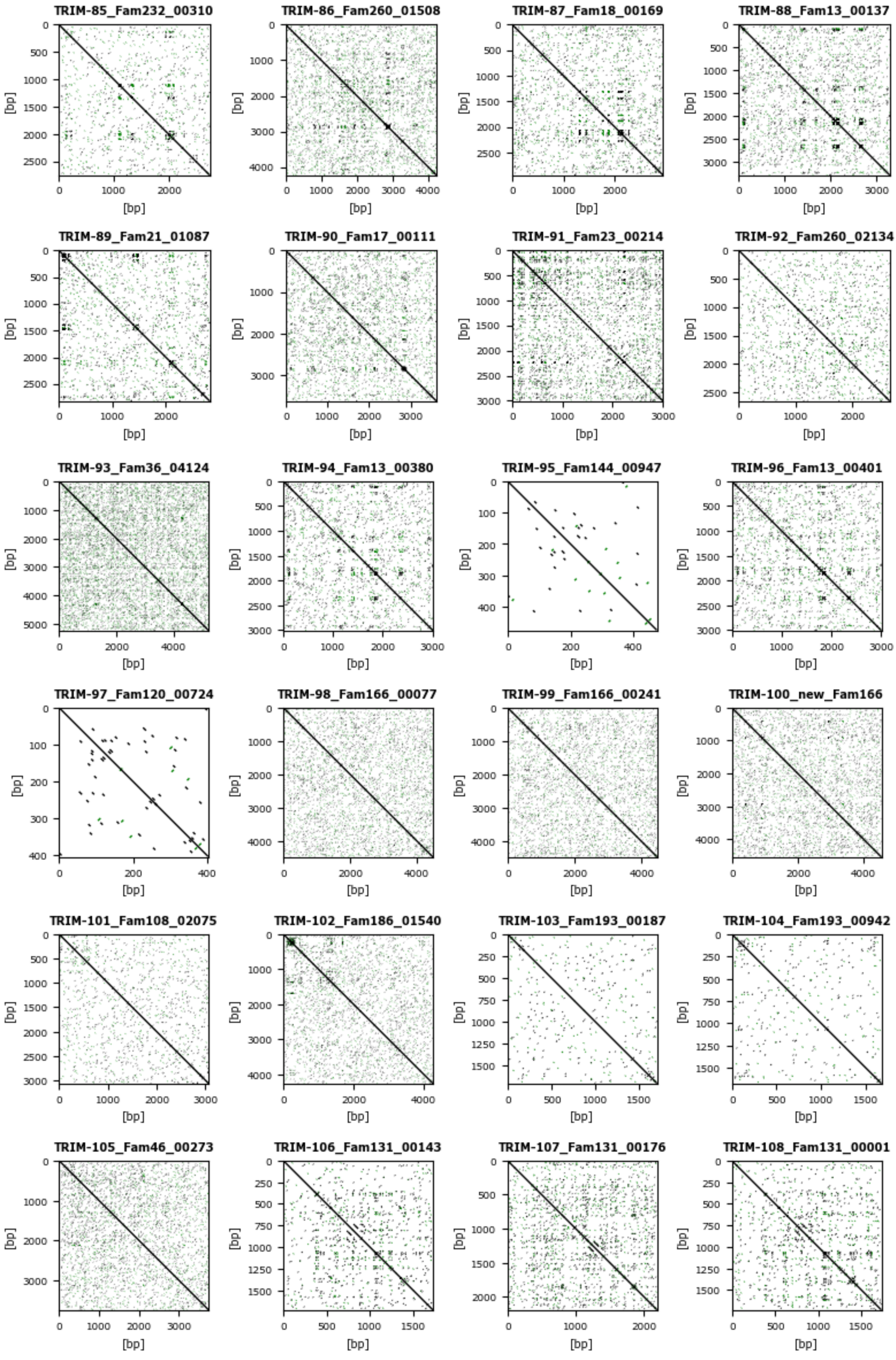

**Additional figure 4: Dotplot vizualisation of TRIM internal sequences.**
Continuation.

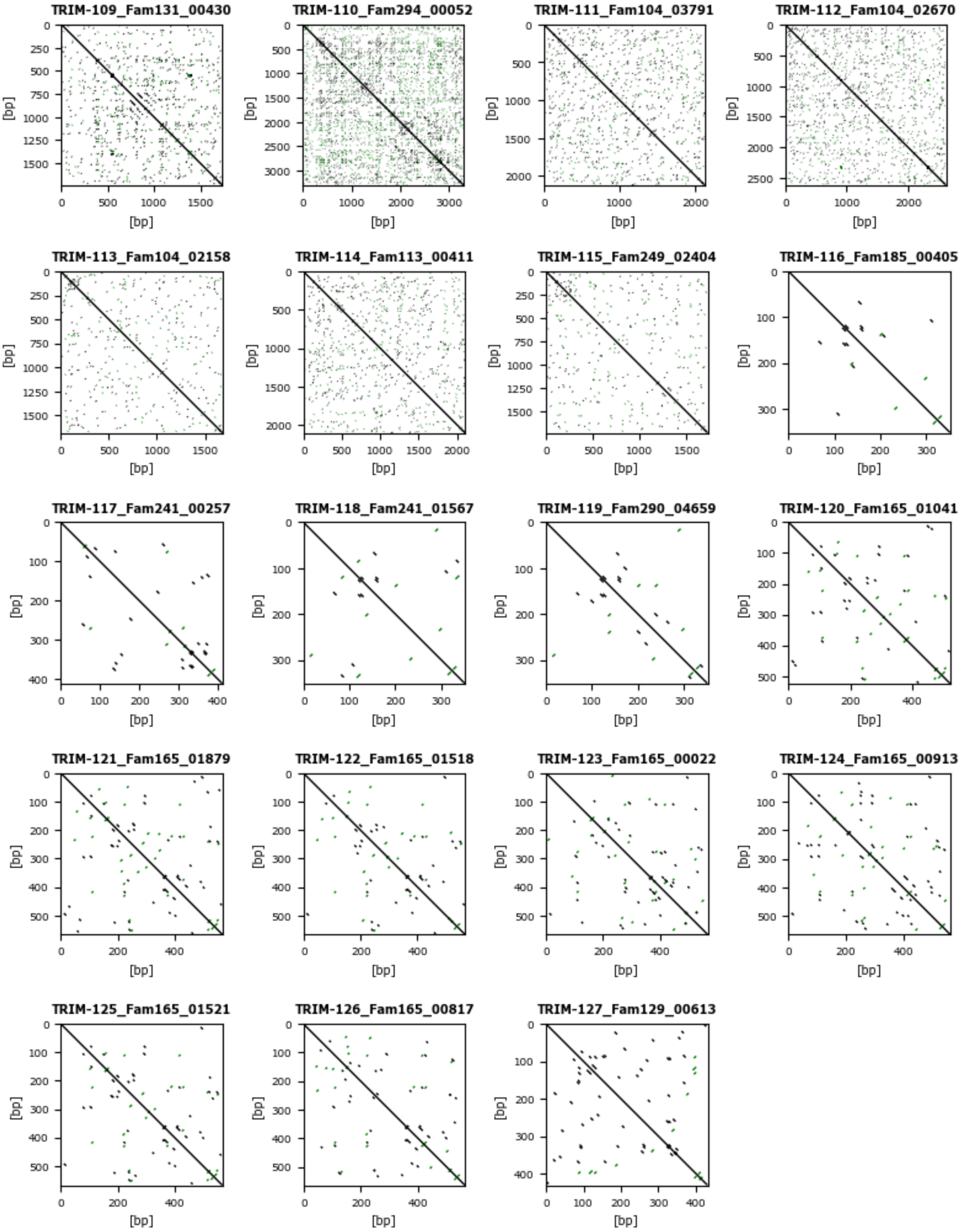

**Additional figure 5: Shared similarity of multiple TRIM families.**

TRIM families are grouped by their modularity and condensed into modularity groups 1-16 as also described in Addtional table 1. For Dotplot vizualisation we used a wordsize of 10 and disabled reverse complement highlighting. Highlighted with blue rectangles are LTR motifs.

**Modularity group M1**

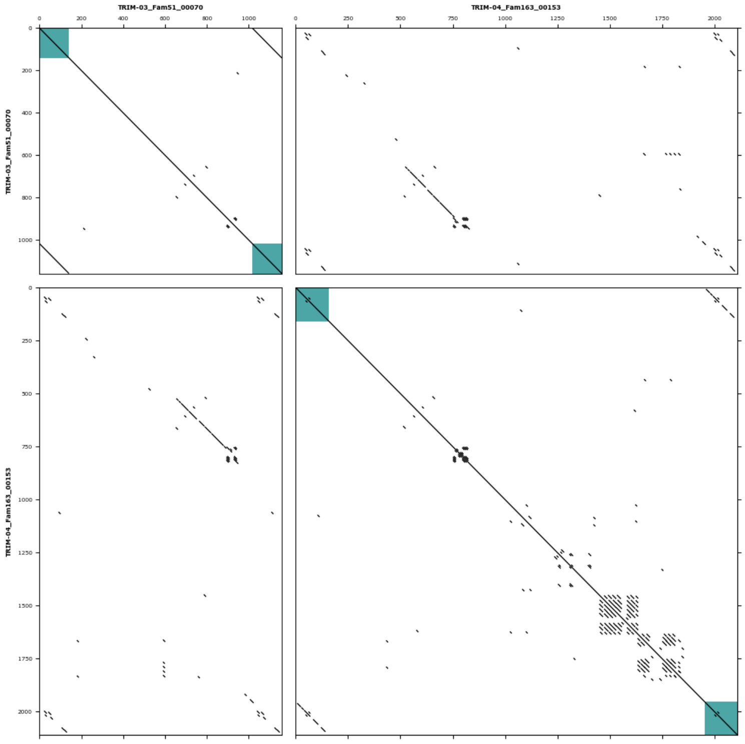


**Modularity group M2**

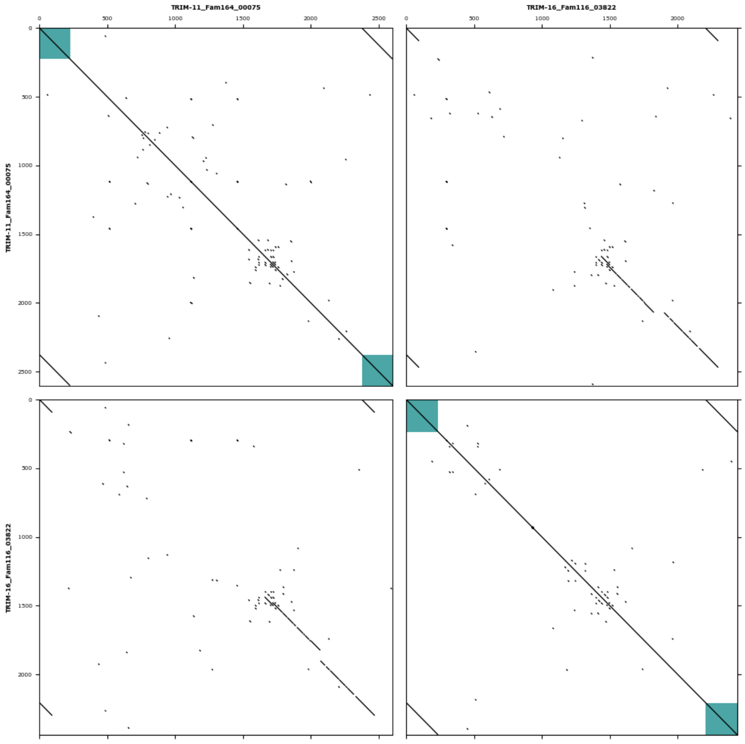


**Modularity group M3**

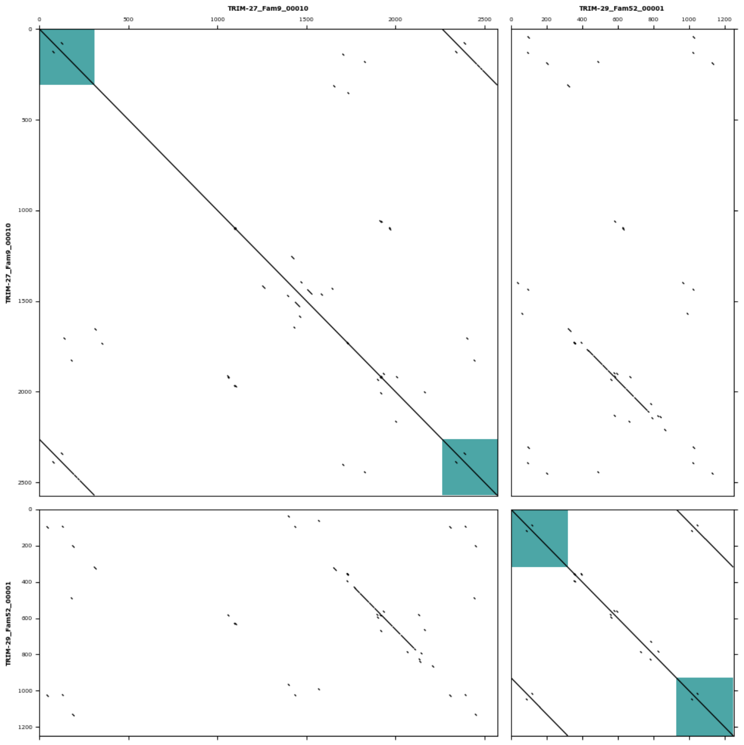


**Modularity group M4**

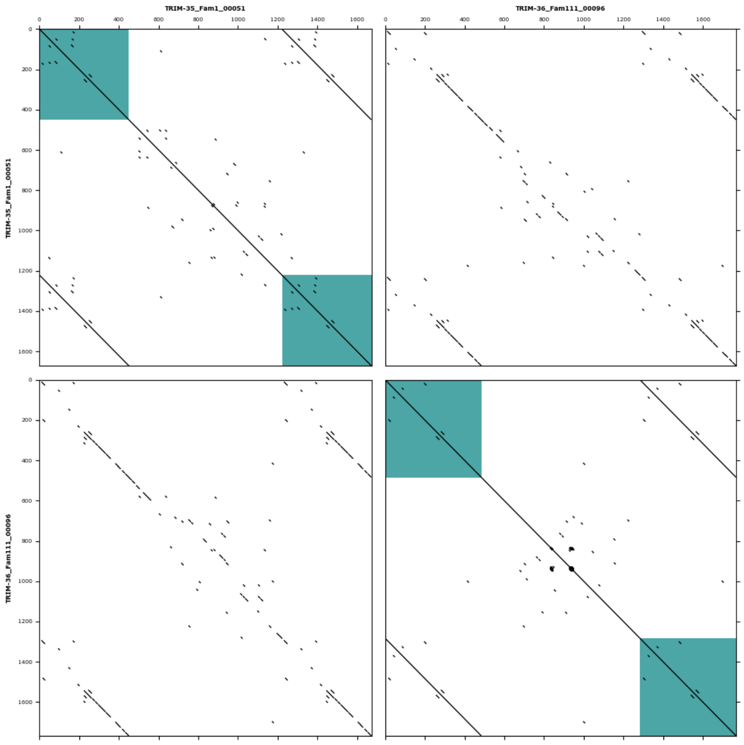

**Additional figure 5: Shared similarity of multiple TRIM families.**
Continuation.

### Modularity group M5

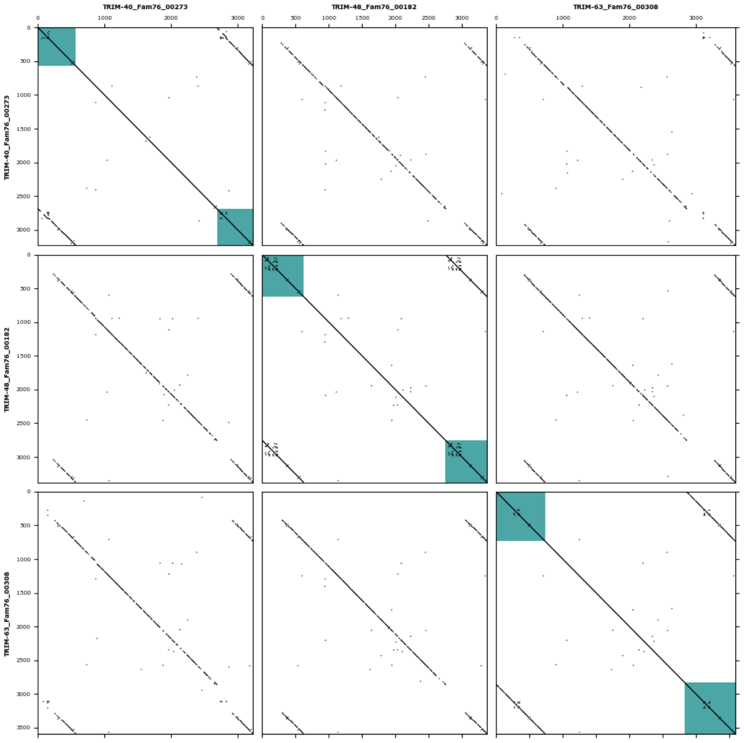


### Modularity group M6

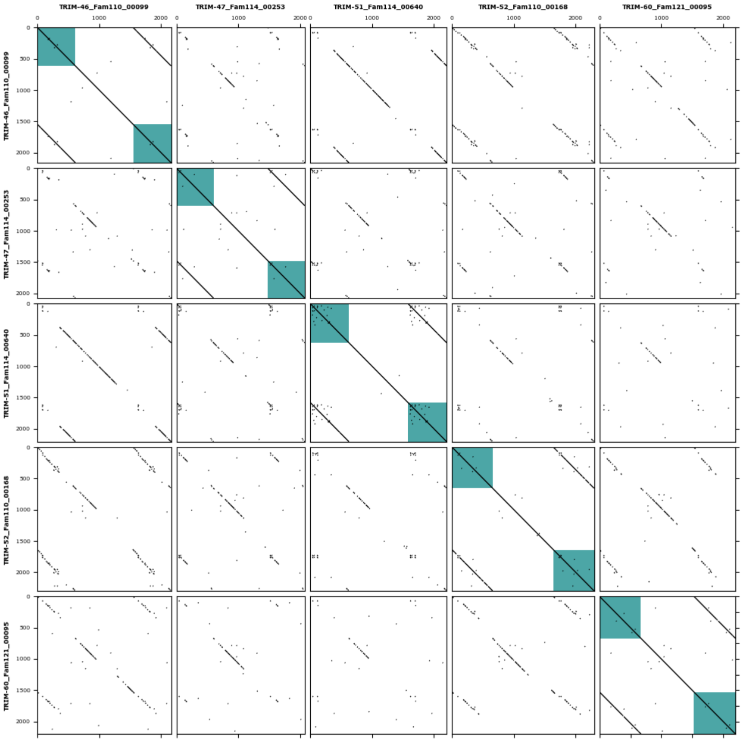


### Modularity group M7

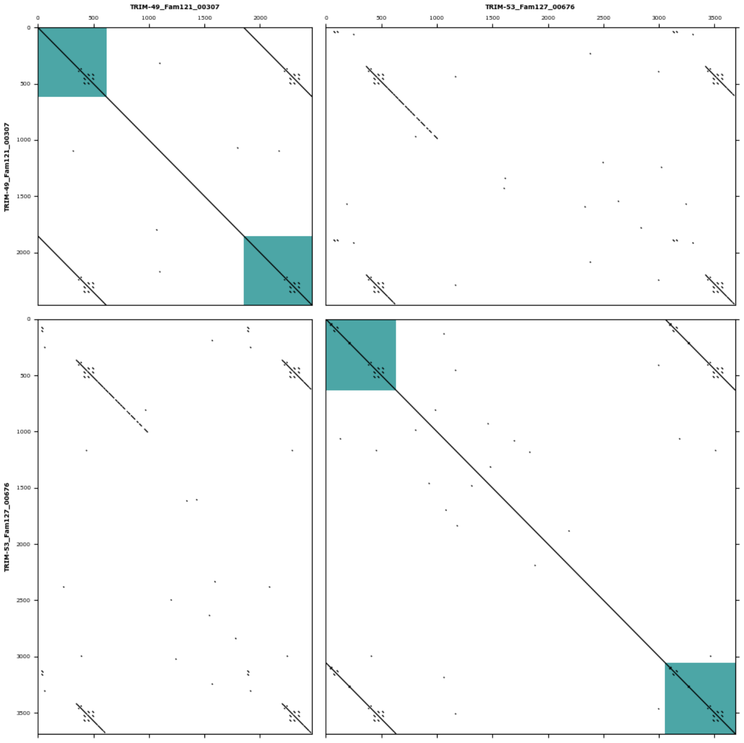


### Modularity group M8

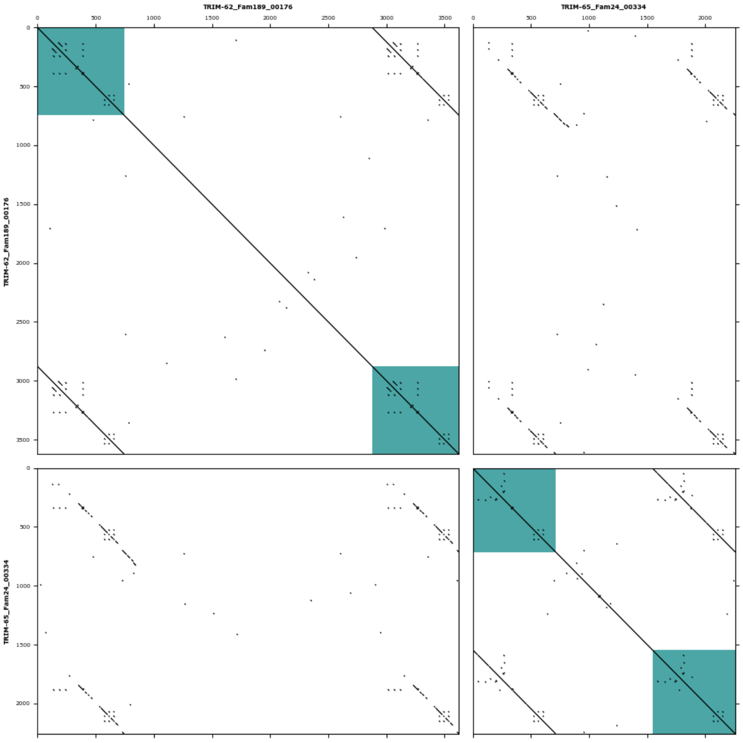

**Additional figure 5: Shared similarity of multiple TRIM families.**
Continuation.

### Modularity group M9

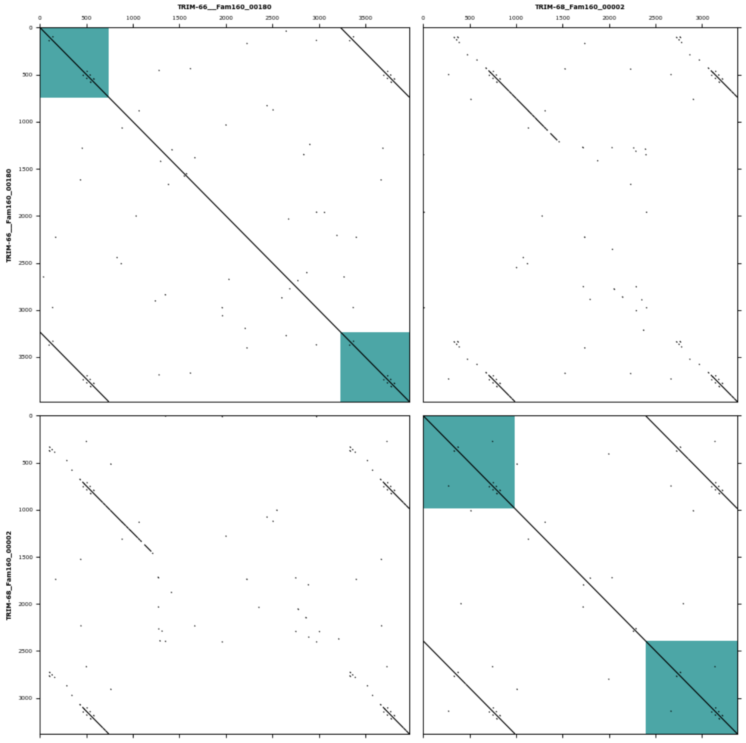


### Modularity group M10

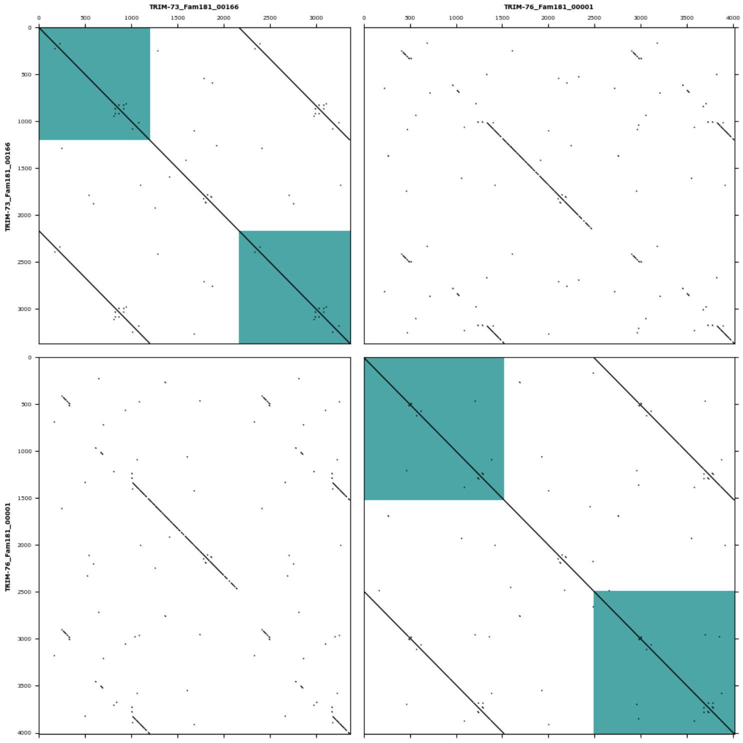


### Modularity group M11

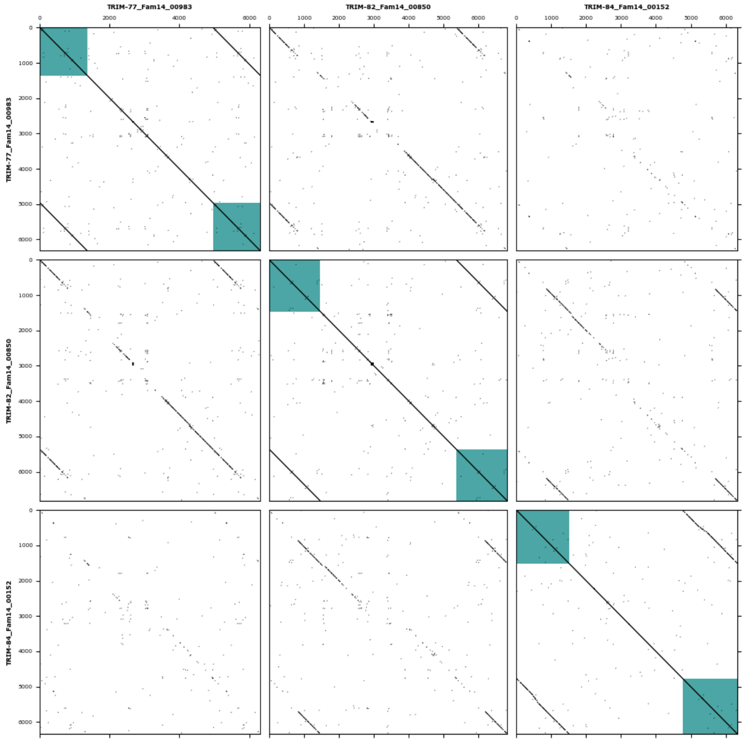


### Modularity group M12

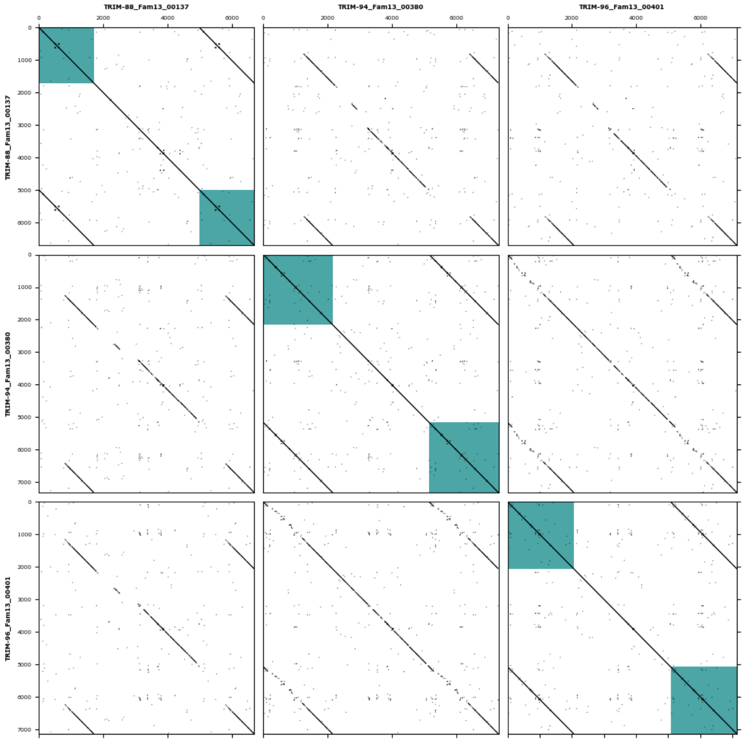

**Additional figure 5: Shared similarity of multiple TRIM families.**
Continuation.

### Modularity group M13

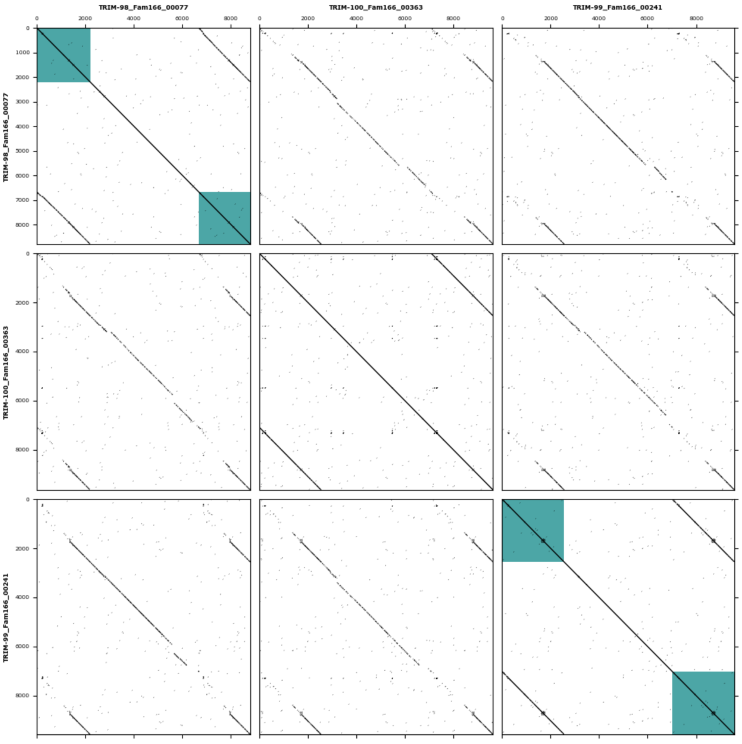


### Modularity group M14

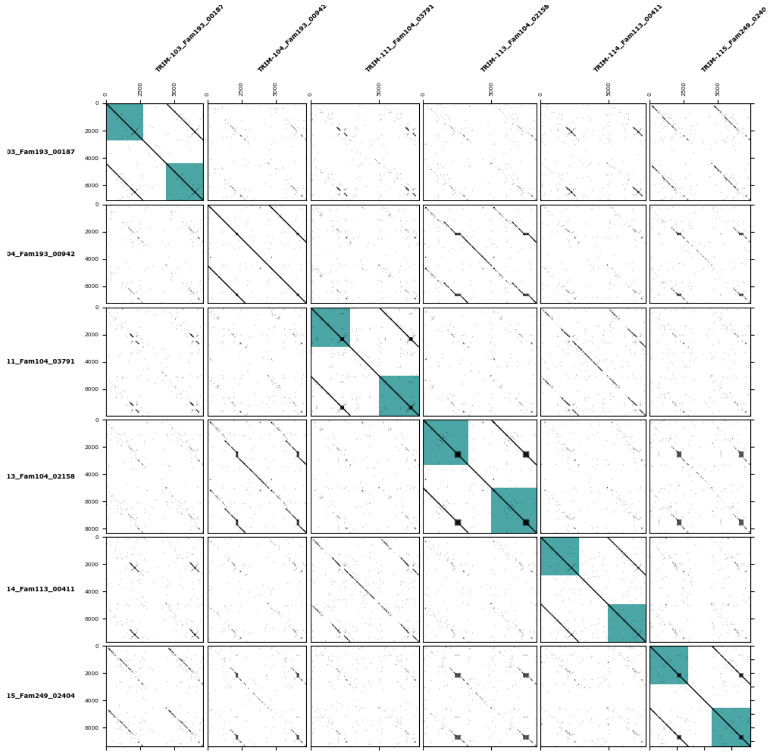

### Modularity group M15

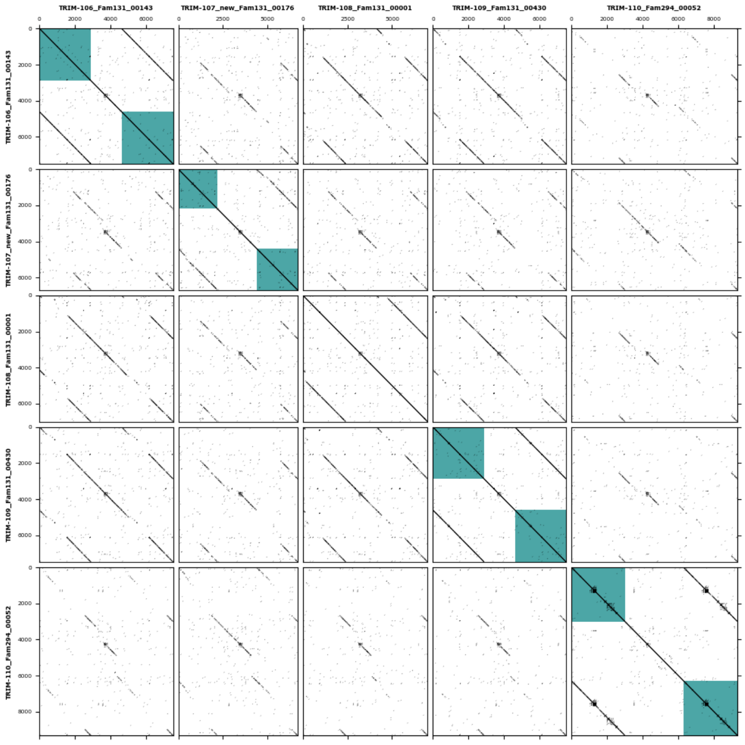


### Modularity group M16

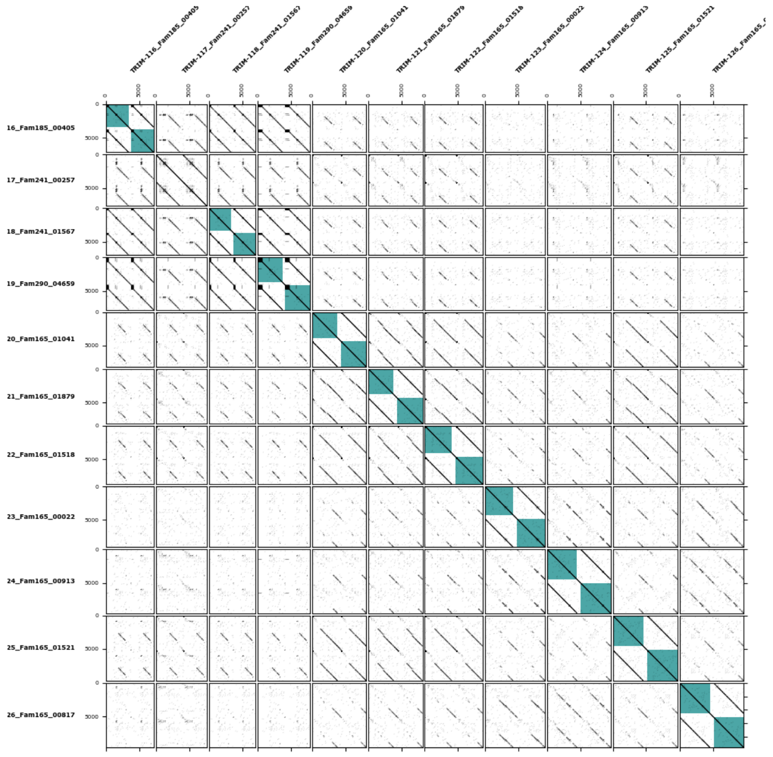
